# Cytosine-to-uracil RNA editing is upregulated by pro-inflammatory stimulation of myeloid cells

**DOI:** 10.1101/2025.03.14.643382

**Authors:** Hyomin Seo, Winston H. Cuddleston, Ting Fu, Elisa Navarro, Madison Parks, Amanda Allan, Anastasia G. Efthymiou, Michael S. Breen, Xinshu Xiao, Towfique Raj, Jack Humphrey

**Affiliations:** Department of Genetics and Genomic Sciences, Icahn School of Medicine at Mount Sinai, New York, NY, USA; Nash Family Department of Neuroscience & Friedman Brain Institute, Icahn School of Medicine at Mount Sinai, New York, NY, USA; Ronald M. Loeb Center for Alzheimer’s Disease, Icahn School of Medicine at Mount Sinai, New York, NY, USA; Icahn Institute for Data Science and Genomic Technology, Icahn School of Medicine at Mount Sinai, New York, NY, USA; Estelle and Daniel Maggin Department of Neurology, Icahn School of Medicine at Mount Sinai, New York, NY, USA; Department of Integrative Biology and Physiology, University of California, Los Angeles, Los Angeles, CA 90095, USA; Instituto Universitario de Investigacion en Neuroquímica, Departamento de Bioquímica y Biologia Molecular, Facultad de Medicina, Universidad Complutense, Madrid, Spain; Centro de Investigacion Biomédica en Red de Enfermedades Neurodegenerativas (CIBERNED), Instituto de Salud Carlos III, Madrid, Spain; Instituto Ramon y Cajal de Investigacion Sanitaria (IRYCIS), Madrid, Spain; Seaver Autism Center for Research and Treatment, Icahn School of Medicine at Mount Sinai, New York, NY, USA; Department of Psychiatry, Icahn School of Medicine at Mount Sinai, New York, NY, USA; Mindich Child Health and Development Institute, Icahn School of Medicine at Mount Sinai, New York, NY, USA; Bioinformatics Interdepartmental Program, University of California, Los Angeles, Los Angeles, CA 90095, USA

## Abstract

Myeloid cells undergo large changes to their gene expression profile in response to inflammatory stimulation. This includes an increase in post-transcriptional modifications carried out by adenosine-to-inosine (A-to-I) and cytosine-to-uracil (C-to-U) RNA editing enzymes. However, the precise RNA editing targets altered by stimulation and the consequences of RNA editing on gene expression and the proteome have been understudied. We present a comprehensive RNA editing analysis of stimulated myeloid cells across three independent cohorts totalling 297 samples, including monocytes and IPS-derived microglia. We observed that C-to-U editing, while less abundant, has a higher effect size in response to stimulation than A-to-I, and has a greater potential to recode the proteome. We investigated the consequences of RNA editing on RNA stability and gene expression using *in silico* and *in vitro* reporter methods, and identified a recoding C-to-U site in *ARSB* that mimics a reported lysosomal storage disorder mutation.

## Introduction

Myeloid cells are the innate immune system lineage which includes monocytes and macrophages in the periphery and microglia in the central nervous system (CNS). Myeloid cells are highly plastic and can change their transcriptional state in response to environmental and disease-related stimuli. Studying the transcriptional regulation of myeloid cells thus has the potential to uncover new insights into their role in inflammatory and neurodegenerative disease. However, while studies have focused on regulatory mechanisms such as chromatin accessibility, gene expression, or alternative splicing^1–6^, comparatively little attention has been paid to the role of RNA modifications.

RNA editing broadly refers to a set of co- and post-transcriptional modifications whereby individual RNA nucleotides are chemically altered by specific enzymes, diversifying the range of RNA molecules which can arise from the same genome sequence. This can lead to a variety of downstream consequences, including recoding of the amino acid sequence, or changes in RNA stability. There are two main types of RNA editing: cytosine-to-uracil (C-to-U) and adenosine-to-inosine (A-to-I). However, because both second-generation sequencing technology and the ribosome read uracil as thymine (T) and inosine as guanine (G), the two categories of RNA editing are also commonly referred to as C-to-T and A-to-G.

The most abundant type of RNA editing is the deamination of A-to-I, catalyzed by the *ADAR* (Adenosine Deaminases Acting on RNA) family of proteins, which target double-stranded RNA substrates, primarily formed by inversely oriented Alu retrotransposable elements^7,8^. In humans, there are three primary ADAR proteins: ADAR1, ADAR2, and ADAR3^9^. For the ADAR1 protein, there is a constitutively expressed isoform (ADAR1p110) and an interferon-inducible isoform (ADAR1p150) which uniquely possesses the ability to shuttle between the nucleus and cytoplasm, while all other ADAR proteins are nuclear localized^10–13^. With the advancement of high-throughput sequencing technologies, millions of A-to-I editing sites have been identified in the human transcriptome^14^. In exons, A-to-I edits have the potential to recode the amino acid sequence of the eventual protein product, and these recoding events are especially appreciated for playing important roles in several aspects of neurophysiology^13^. However, in line with the distribution of Alu elements throughout the human genome, A-to-I edits primarily reside in non-coding regions^15^, where these events have been demonstrated to impact various aspects of gene regulation including RNA splicing and stability^16–18^.

The deamination of C-to-U is catalyzed by the APOBEC (Apolipoprotein B mRNA Editing Catalytic Polypeptide-like) family of proteins, comprised of 11 primary isoforms in humans, which target both DNA and RNA^19^. APOBEC3A and APOBEC3G have been identified to act as editing enzymes in human immune cells^20–22^. While less frequent than A-to-I editing, C-to-U editing can significantly impact RNA structure, stability, splicing, and translation, ultimately affecting gene expression and protein function^23^. For example, C-to-U editing of the mitochondrial complex protein SDHB creates an in-frame stop codon, reducing protein levels^13,24^. C-to-U editing events occur more frequently in exonic regions than A-to-I^20–22^, suggesting that these events have a higher recoding potential.

Moreover, RNA editing can be activated by external stimuli, including inflammatory responses, where C-to-U edits have been shown to contribute to immune-related gene expression changes essential for effective pathogen defense^20–22^. Both A-to-I and C-to-U RNA editing are induced in response to stimuli like hypoxia and interferons, influencing differentiation and function of immune cells, including monocytes and macrophages^25–27^.

In this study, we analyze blood-derived monocyte samples stimulated with either lipopolysaccharide (LPS) or interferons (IFN) to investigate the interplay between RNA editing and gene expression dynamics, and replicate our findings in independent cohorts of monocytes and induced pluripotent stem cell-derived microglia (iMicroglia). We further explore the functional consequences of stimulation-induced RNA editing with both *in silico* and *in vitro* reporter assays. Stimulating myeloid cells with IFN and LPS has been established as an experimental model for the immune activation observed in neurodegenerative disease contexts^28–31^. This work therefore lays a foundation for understanding the role of RNA editing in both homeostatic and disease processes within myeloid cells.

## Results

### Pro-inflammatory stimulation induces RNA editing in monocytes

This study incorporates three distinct cohorts of stimulated myeloid cells. Our discovery cohort consists of monocytes purified from peripheral blood (n=165) and was generated for an ongoing study. The discovery cohort contains 55 unique donors stimulated with lipopolysaccharide (LPS) and interferon beta (IFNβ). **(Figure 1A).** To validate and extend the findings from the discovery cohort, we used a previously generated^29^ cohort (n=105), consisting of 35 unique donors stimulated with LPS or interferon-gamma (IFN*γ*) to serve as a replication **(Figure 1B)**. The two cohorts were generated with different RNA-seq library preparations, resulting in noticeable differences in coverage over exons and introns. To assess whether changes in RNA editing observed in monocytes can be identified in microglia, we then compared RNA editing in a third cohort composed of 27 stimulated induced pluripotent stem cell-derived microglia (iMicroglia) samples derived from multiple differentiations of a single donor line^29^ and stimulated with LPS or IFN*γ* **(Figure 1C).** Our systematic design allows for comprehensive analyses across different stimulation conditions and cell types.

**Figure 1.**
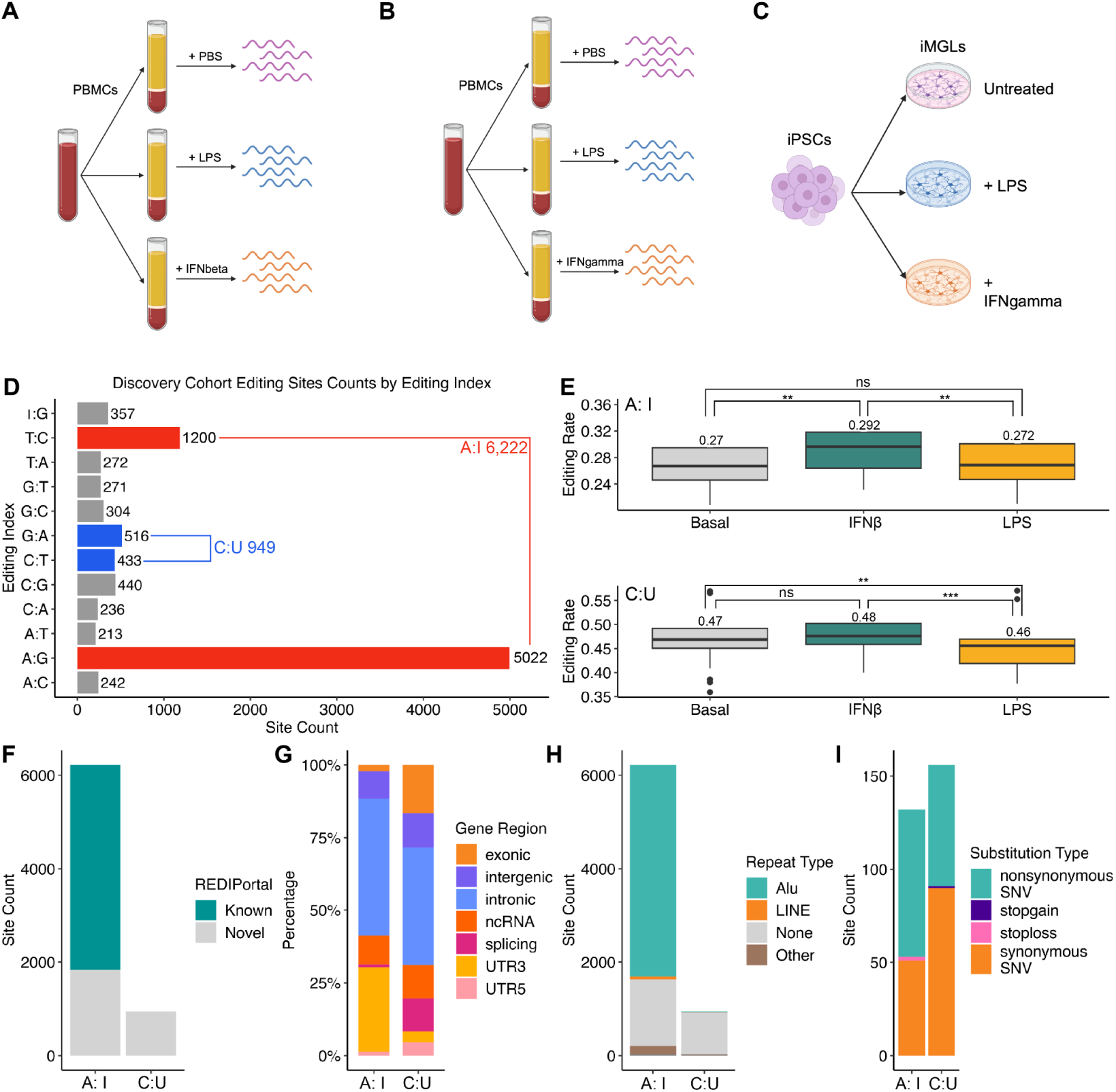
RNA editing sites within the discovery cohort exhibit varying editing rates depending on the stimulation status. Design of the study, including schematics for the discovery cohort **(A)**, replication cohort **(B)**, and iMicroglia cohort **(C)**. **D)** Distribution of editing sites found in the discovery cohort for 12 different editing indexes. **E)** Mean editing mean rates for A-to-I editing sites (upper panel) and C-to-U sites (lower panel) split by each stimulation condition. P-values from paired t-test. ns: non significant; **: P < 0.01; ***: P < 0.001. Counts of editing sites stratified by presence in the REDIportal database (**F**), genomic region (**G**), RepeatMasker annotations (**H**), and predicted substitution type for all A-to-I and C-to-U sites (**I**). LINE; Long interspersed nuclear element, SNV; single nucleotide variant.

We processed each cohort through a comprehensive RNA editing detection pipeline which quantifies editing rates at both known A-to-I editing sites from the REDIportal database^32^ and discovers novel editing sites, which includes A-to-I and C-to-U sites. Sites were filtered both per sample and across samples and stimulation conditions with strict filters to ensure robustness. In the discovery cohort, we identified a total of 9,506 editing sites across the 12 possible substitution types (A:C, A:G, A:T, C:A, C:G, C:T, G:A, G:C, G:T, T:A, T:C, and T:G). Among them, 6,222 potential A-to-I (combining A:G and the reverse complement T:C) (65.45%) and 949 C-to-U (combining C:T and the reverse complement G:A; 9.98%) editing events were found, suggesting that 75% of the observed RNA substitutions are potential RNA editing events **(Fig. 1D; Supplementary Table 1).** Taking the average editing rate within each stimulation group, we observed that editing rates for both types were elevated by IFNβ stimulation compared to baseline. **(Fig. 1E; Supplementary Table 1).** Although fewer C-to-U editing sites were detected compared to A-to-I sites, the magnitude of change in C-to-U rates was as large, if not larger, than that of A-to-I sites. This trend is observed in editing events detected under both LPS and IFNβ stimulation **(Supplementary Fig. 2A-B; Supplementary Table 1)**. 65.9% of the A-to-I sites have been previously reported in the REDIPortal database (v2.0)^32^ (**Fig. 1F; Supplementary Table 2)**. Annotations for editing events in the A-to-I and C-to-U editing indexes revealed distinct distributions across genomic regions. As expected, the majority of A-to-I sites were located in intronic (47.18%) and 3’UTR regions (29.05%), with a minority located within exons (2.15%) **(Fig. 1G; Supplementary Table 2)**. The majority (72.83%) of intronic A-to-I editing events were found to overlap with *Alu* elements **(Fig. 1H; Supplementary Table 2)** Conversely, 157 C-to-U sites (16.54%) were located within exons **(Fig. 1G; Supplementary Table 2)**. Of the exonic sites, 79 A-to-I and 65 C-to-U sites were predicted to cause non-synonymous single nucleotide variants (SNVs) and act as potential recoding sites **(Fig. 1I; Supplementary Table 2).** To determine whether the change in editing rates due to stimulation is specific to A-to-I and C-to-U events, we analyzed the frequency of all other possible nucleotide substitutions. We found that, unlike A-to-I and C-to-U events, the “editing rates” for all other types either remained unchanged or decreased in response to stimulation, suggesting these substitution types are likely background noise **(Supplementary Fig. 1A; Supplementary Table 22).**

Leveraging the large size of our cohort, we then assessed the variance in RNA editing both globally and at each site explained by a set of clinical and technical variables. Using principal component analysis, we observed that the first principal component strongly correlated with stimulation status and the levels of *ADAR* and *APOBEC* gene expression, which are highly upregulated in both stimulation conditions (**Supplementary Fig. 1B; Supplementary Table 23**).

The second principal component correlated with technical metrics derived from read coverage over exons and introns (**Supplementary Fig. 1C,F; Supplementary Table 24)**. Assessing the variance explained in each site, we observed that *APOBEC3G* and *APOBEC3A* expression explained the most variation per site on average, followed by read coverage and batch for both indexes **(Supplementary Fig. 1D,G; Supplementary Table 24).** We also observed the principal components influenced by the stimulation status of each sample and the expression of *APOBEC3* genes, both for A-to-I and C-to-U editing rates. Notably, samples stimulated with IFNβ were visibly separated from other samples in principal component space. These IFNβ stimulated samples also exhibited higher expression levels of the *APOBEC3A* gene and *APOBEC3G* gene **(Supplementary Fig 1E, H; Supplementary Table 24).**

We performed a differential editing analysis on the discovery cohort samples, comparing the effect of LPS or IFNβ stimulation with baseline samples, correcting for technical (ribosomal base coverage percentage) and clinical (diagnosis, age, sex) covariates. A total of 7,171 editing sites were tested for LPS stimulation, revealing 1,228 differentially edited sites (DESs) at FDR < 0.05, accounting for 17.12% of all tested sites. Of these, 1,037 were A-to-I and 191 were C-to-U **(Fig. 2A; Supplementary Table 3)**. 505 A-to-I editing events (48.69%) were in the 3’ UTR whereas 282 were within introns. The majority (96.62%) of A-to-I events occurred in non-coding regions. 74 C-to-U differentially edited events were within introns, and 45 were within 3’ UTRs. 13 of the exonic C-to-U editing sites were predicted to cause nonsynonymous SNV mutations **(Supplementary Table 2)**. One site in *APOBEC3F* was predicted to lead to a stopgain mutation **(Supplementary Table 2)**.

**Figure 2.**
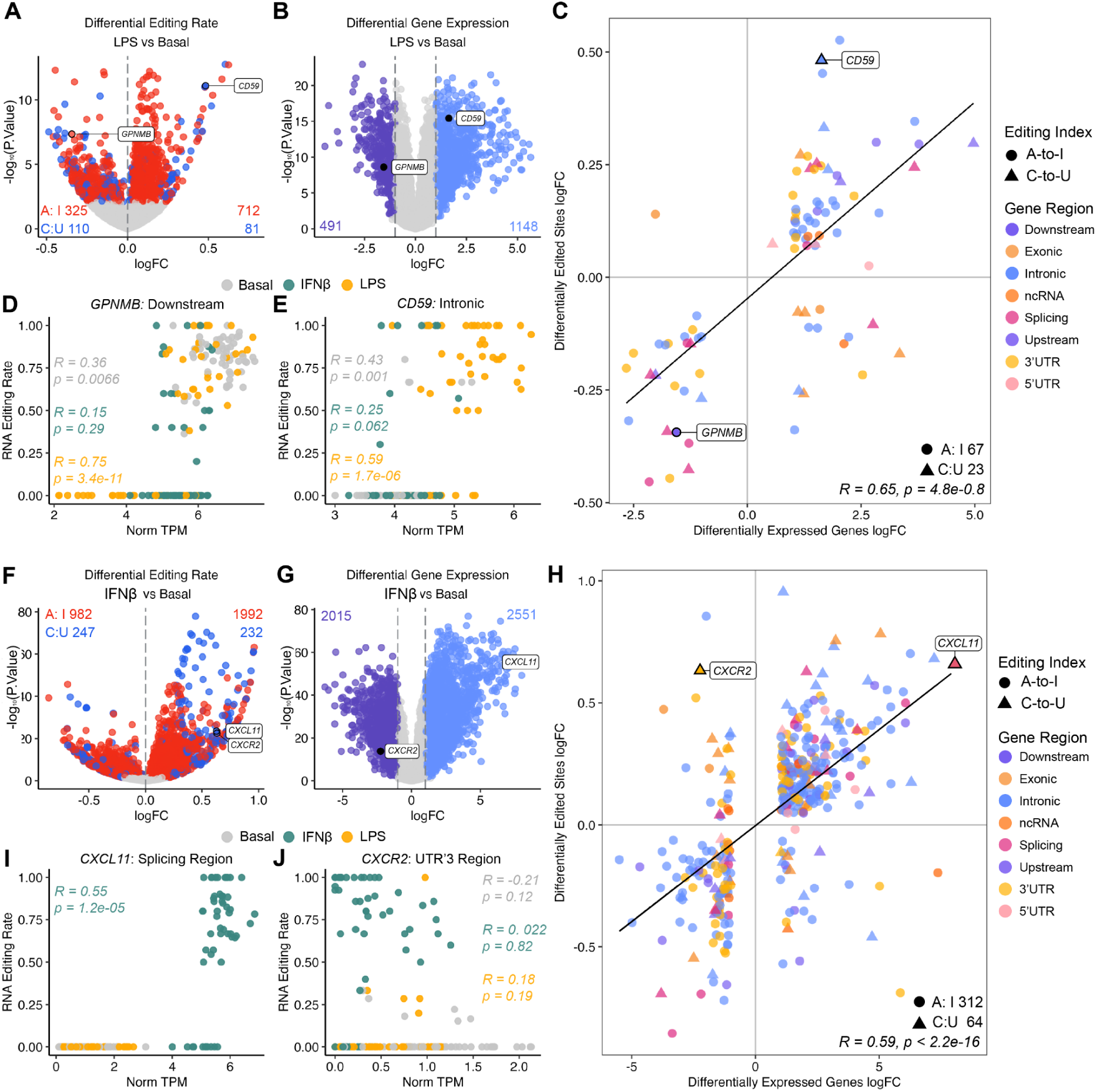
Differential RNA editing and gene expression in the discovery cohort. **A)** Differential editing analysis comparing LPS-stimulated samples and basal samples. **B)** Differential gene expression analysis comparing LPS-stimulated samples and basal samples. **C)** Comparing LogFC of DEGs overlapping DESs, colored by gene location. **D)** Gene expression as normalized TPM plotted against RNA editing rates of *GPNMB*, with an A-to-I editing site located in the downstream region, chr7:23275201. **E)** *CD59,* with a C-to-U editing site located within an intron, at chr11:33717481. **F)** Differential editing analysis between IFNβ-stimulated and basal samples. **G)** Differential gene expression analysis between IFNβ-stimulated and basal samples. **H)** Comparison of DEGs that overlap DESs, colored by gene location. **I)** Gene expression plotted against RNA editing rates of *CXCL11*. **(J)** Gene expression plotted against RNA editing rates of *CXCR2*.

In IFNβ stimulation, 2,974 A-to-I and 479 C-to-U DES were detected to be significantly edited, with C-to-U sites having notably higher effect sizes (log_2_ fold-changes; logFC) than A-to-I **(Fig. 2F; Supplementary Table 6)**. Among the A-to-I DESs, 1,159 sites were within 3’UTRs, and 1,128 sites were within introns. 46 A-to-I sites were predicted to cause a nonsynonymous SNV and one site as a potential stop-loss mutation. For the 479 C-to-U DESs, 193 sites were within introns, and 109 were within exons. 42 C-to-U sites were predicted to cause nonsynonymous SNVs, while one stopgain site was detected **(Supplementary Table 2)**. The site in *APOBEC3F* detected with a stopgain mutation, chr22:39052279, was also identified under LPS stimulation. However, this site exhibited a greater decrease in editing under IFNβ stimulation (logFC = -0.155) compared to LPS stimulation (logFC = -0.082). While *APOBEC3F* has not been identified to have cytidine deaminase activity, we propose that changes in its expression caused by C-to-U editing could act to regulate C-to-U editing through its family members. Additionally, we examined the effect sizes of 4,011 A-to-I and 670 C-to-U sites observed under both stimuli **(Supplementary Fig. 2C; Supplementary Table 3, 6)**. While A-to-I events showed significant differences in logFC between IFNβ and LPS, no such difference was observed for C-to-U events. To assess the impact of *ADAR* and *APOBEC* gene expression as covariates in differential editing analysis, we repeated the analysis using a second model that included the expression of *APOBEC3G, APOBEC3F,* and *APOBEC3A* as covariates. This resulted in a significantly lower number of detected edited sites: 505 sites under LPS and 188 sites under IFNβ stimulation. These findings confirm the direct role of *ADAR* and *APOBEC* genes in regulating RNA editing, as supported by previous studies **(Supplementary Fig. 2D, Supplementary Tables 25, 26).**

In a previous study examining RNA editing on A-to-I and C-to-U indexes under hypoxia treatment and macrophage polarization conditions, 3,304 editing events were identified^21^. When comparing the reported sites to the 30,957 sites detected at least once across our three cohorts, we identified 171 overlapping A-to-I and 17 C-to-U sites between the studies. In total, 523 unique genes were found to contain editing events in both studies, with 111 genes sharing 188 exact overlapping sites. This comparison was based on supplementary data obtained directly from the paper, lifted over from GRCh37 to GRCh38.

### RNA editing changes are largely concordant with gene expression changes

We performed differential gene expression analysis for samples in our discovery cohort to compare the overlap at the gene level between differentially expressed genes (DEGs) and our DES. The majority of genes in the *ADAR* and *APOBEC* gene families were upregulated under either LPS or IFNβ stimulation (**Supplementary Fig. 1B; Supplementary Table 23**). Furthermore, including *ADAR* and *APOBEC* gene expression as additional covariates for differential editing significantly reduced the DES discovery rate, suggesting that the editing changes observed in stimulation are largely mediated by changes in the expression of the editing enzymes **(Supplementary Fig. 1C,F)**. Under LPS stimulation, 1,148 up-regulated DEGs and 491 down-regulated DEGs were identified **(Fig. 2B; Supplementary Table 4)**. Notably, DEGs displayed large changes in expression, ranging from -4.4 to +5.6, indicating changes from an 8-fold decrease to a 32-fold increase.

We conducted a comparative analysis of the DESs and DEGs identified in each stimulation to investigate the potential correlation between RNA editing rate and gene expression. Under LPS stimulation, we identified 90 unique DEGs (5.5% of the total DEGs) associated with 255 unique DESs (20.7% of the total; 208 A-to-I and 46 C-to-U sites). Therefore, RNA editing changes were found in only a minority of genes differentially expressed by LPS stimulation, whereas a larger proportion of DES also have an accompanying DEG. Each gene was then paired with an editing site that had the largest absolute DES logFC value, leaving 90 unique genes associated with 90 editing sites to examine the concordance of effect direction. The gene region of each 90 sites and their respective editing index are identified **(Fig. 2C)**. We demonstrated that sites have largely concordant directions of effect between editing and expression. We observed that sites that are increased in editing tend to correspond to up regulated gene expression at the same loci and *vice versa* (R = 0.65, *p* = 4.8e-0.8) **(Fig. 2C).**

In the gene *GPNMB*, previously linked to Parkinson’s disease through GWAS^33^, we identified an exonic A-to-I site located in the downstream region (chr7:23275201), likely an unannotated 3’UTR. This site showed a notable concordance in the direction of effect between RNA editing and gene expression under stimulation. Particularly in LPS-stimulated samples, the high editing rate was associated with high expression **(Fig. 2D; Supplementary Table 5)**. This finding aligns with previous studies showing that *GPNMB* expression is increased under pro-inflammatory conditions like LPS treatment, and in pathological conditions, such as neurodegenerative diseases and cancer^34^. Additionally, a C-to-U editing site located in the intronic region of *CD59* was identified (chr11:33717481), with LPS-stimulated samples displaying both increased editing rate and increased expression, while basal and IFNβ-stimulated samples were mostly unaffected **(Fig. 2E; Supplementary Table 5).**

Gene expression under IFNβ stimulation revealed 2,551 up-regulated genes and 2,015 down-regulated genes **(Fig. 2G; Supplementary Table 7)**. Under IFNβ stimulation, we detected 376 DEGs linked to 1,376 unique DESs (1,200 A-to-I and 176 C-to-U sites; 8.2% of the total DEGs; 39.8% of total DES). We matched each gene with the editing site that had the largest absolute DES logFC, resulting in 376 DEGs associated with 376 editing sites and compared their logFC of DES and DEG analysis **(Fig. 2H)**, also identifying largely concordant directions of effect, with some outliers. Notably, *CXCL11* was identified as highly upregulated in both RNA editing and gene expression. *CXCL11* was associated with an exonic C-to-U editing site (chr4:76034817). Interestingly, RNA editing at this site showed a marked difference between IFNβ-stimulated samples and the controls **(Fig. 2I; Supplementary Table 5)**. It was previously studied that chemokines like *CXCL11* and similar chemokines are significantly upregulated in response to inflammatory stimuli like IFNβ and are involved in immune responses of the CNS^35^. Conversely, *CXCR2*, with a C-to-U editing site in the 3’ UTR (chr2:218135891), showed a discordant relationship, with an upregulated RNA editing rate but downregulated expression, particularly in IFNβ-stimulated samples **(Fig. 2J; Supplementary Table 5)**. Previously, it was demonstrated that in a viral model of neuroinflammation and demyelination, genetic silencing of *CXCR2* increased remyelination^36^.

### Differential editing replicates in an independent monocyte cohort

In our replication cohort of monocytes (n=105; 35 samples per group)^29^, a total of 20,552 editing sites were identified, encompassing 20,233 A-to-I and 319 C-to-U sites (**Fig. 3A; Supplementary Table 8)**. Notably, our replication cohort exhibited almost three times more A-to-I editing sites as our discovery cohort, but ⅓ fewer C-to-U sites. We investigated the causes of this discrepancy. The choice of RNA extraction and selection kits in RNA-seq libraries can cause large differences in the coverage of exons and introns, which impacts the ability to quantify RNA editing in these genomic features **(Supplementary Fig. 3A)**. We observed that the discovery cohort was enriched for exonic coverage, whereas the replication cohort exhibited proportionally higher intronic coverage. The A-to-I discovery rate (sites per million mapped bases) negatively correlated with % coding base coverage in the discovery cohort, in line with the location of most A-to-I sites within introns **(Supplementary Fig. 3B)**. However, the replication cohort exhibited a positive correlation between A-to-I discovery and % coding base coverage, suggesting a more complex cause of the discrepancy. Indicative of the disparity in the discovery rates of editing events, the replication rates between the cohorts differed substantially between the two editing types. Specifically, of the 6,022 A-to-I sites found in the discovery cohort, 3,030 (50.3%) could be identified in the replication cohort, whereas of the 949 C-to-U sites, only 102 (10.7%) were replicated.

**Figure 3.**
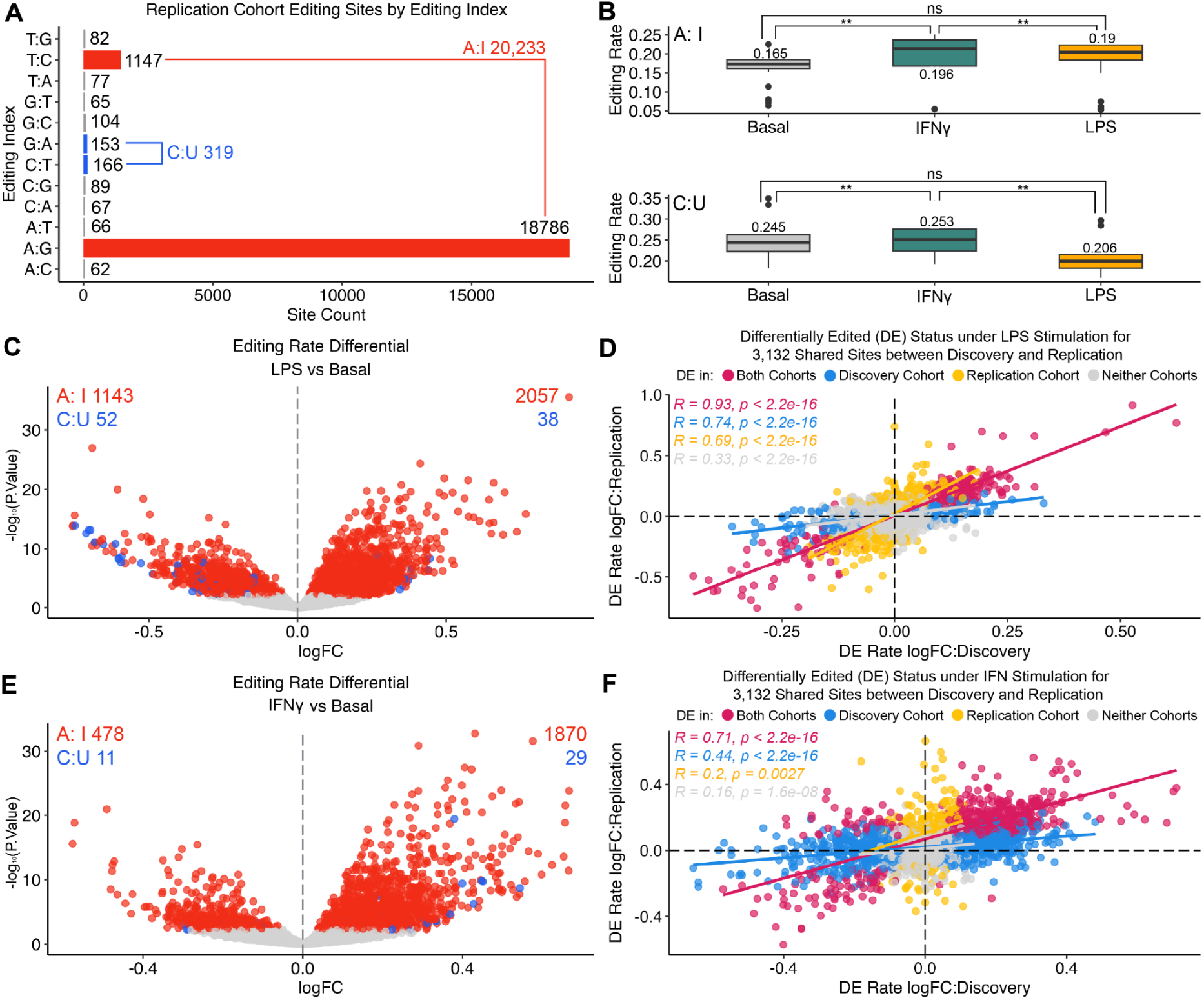
Differential RNA editing identified in the replication cohort. **A)** All editing sites identified in the monocyte replication cohort (n=105). **B)** Distribution of RNA editing rates by editing indexes and stimulation status. P-values from two-sided paired t-test. ns: non significant , P < 0.05; **. **C)** RNA Editing rate differential analysis comparing LPS-stimulated and basal samples. 1,143 A-to-I and 52 C-to-U down-regulated sites and 2,057 A-to-I and 38 C-to-U up-regulated sites were identified (FDR<5%). **D)** Comparing differential editing LogFC identified under LPS stimulation within the discovery cohort and the replication cohort plotted for 3,132 replicated editing sites. Correlations performed using Spearman correlation test. **E)** As in **(C)**, but comparing IFN*γ*-simulated and basal samples. **F)** As in **D)**, but under IFN*γ*.

Another key technical covariate in the replication cohort was the batch in which samples were processed. The cohort was divided into two batches, and we found significant differences in coverage between them. Specifically, the percentage coverage of intronic bases, ribosomal bases, and RNA integrity number (RIN) values varied significantly between batch 1 and batch 2 **(Supplementary Fig. 4A, Supplementary Table 27).** To account for batch variability, batch was included as a covariate in the differential analysis model for the replication cohort. Consistent with the discovery cohort, the expression of *ADAR* and *APOBEC* family genes was notably higher in IFN-stimulated samples **(Supplementary Fig. 4B, Supplementary Table 28).**

The overall mean editing rate for A-to-I editing sites was 0.183, while that for C-to-U sites was 0.234. In both editing indexes, the mean editing rate was highest for IFNγ-stimulated samples, with rates of 0.196 for A-to-I editing sites and 0.253 for C-to-U editing sites **(Fig. 3B; Table 8)**. We performed differential editing analysis in the replication cohort, adjusting for the same covariates shown to be influential to the editing rate (batch, sex, age, % ribosomal base coverage) **(Supplementary Fig. 4C,F; Supplementary Table 27)**. In comparing the effect of LPS, 3,290 DESs were found, consisting of 3,200 A-to-I and 90 C-to-U editing events **(Fig. 3C; Supplementary Table 9)**. In comparing the effect of IFNγ, 2,348 A-to-I and 40 C-to-U DESs were identified **(Figure 3E; Supplementary Table 10)**. When comparing the logFC of the 3,132 editing sites found in both discovery and replication cohorts, we observed a high correlation of both direction and strength of effect, even in sites only called significant in one cohort (both cohorts: R = 0.71, p < 2.2e-16, discovery cohort only : R = 0.44, p < 2.2e-16, Replication cohort only : R = 0.2, p = 0.0027) **(Figure 3D, F)**. Altogether this suggests high replication of DESs between the two cohorts, although differences in detection power attenuate the calling of significant DES in both cohorts.

### Differential editing in IPS-derived microglia

We next assessed whether the effects of stimulation on monocytes could also be seen in microglia, the myeloid cells of the nervous system. We first examined the expression of *ADAR* and *APOBEC* family genes using an independent cohort of IPS-derived microglia (iMicroglia; n=27; 9 samples per group). Consistent with findings from other cohorts, samples stimulated with IFN exhibited the highest expression levels for *ADAR* and *APOBEC* family genes **(Supplementary Fig. 5A; Supplementary Table 29).** Subsequent analyses revealed that the variance in editing rates was significantly influenced by the expression of *APOBEC3G, APOBEC3F, ADAR*, and *ADARB1* genes, along with several technical variables **(Supplementary Fig. 5B-E; Supplementary Table 29, 30).**

We identified 13,329 A-to-I and 1,085 C-to-U sites respectively in iMicroglia **(Fig. 4A; Supplementary Table 11)**. Stimulated samples exhibited a higher global A-to-I editing rate compared to basal samples, whereas the global C-to-U editing rate was only significantly increased in LPS stimulation **(Fig. 4B)**. We performed a differential analysis on editing rates, identifying a significant number of differentially edited sites under both LPS (811 A-to-I; 20 C-to-U) and IFN*γ* (2445 A-to-I; 81 C-to-U) stimulation **(Fig. 4C,E; Supplementary Tables 12, 13).** The majority of the DESs were upregulated, with higher logFC observed for C-to-U sites. A total of 14,414 editing sites were identified in both the discovery and iMicroglia cohorts. When comparing the logFC values between cohorts, no significant correlation was observed for sites detected under LPS stimulation (R=0.092, *p = 0.25*) **(Fig. 4D)**. However, a positive correlation was found for shared sites identified under IFNγ stimulation (R=0.36, *p < 2.2e-16*) **(Fig. 4F).** Therefore, we suggest that stimulation-induced RNA editing changes can be identified in microglia-like cells, as well as monocytes.

**Figure 4.**
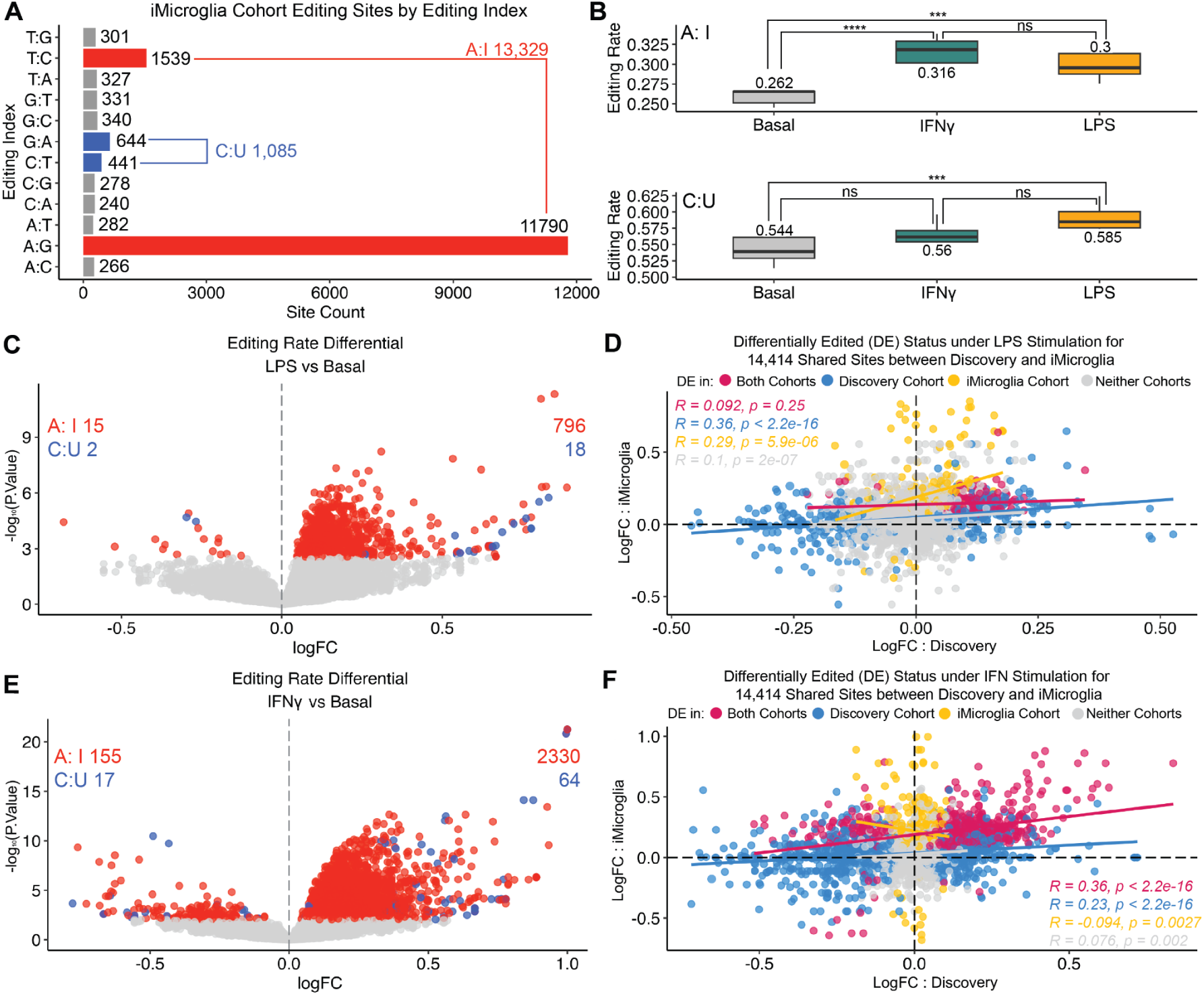
Differentially edited sites and expressed genes identified in IPS-derived microglia. **A)** RNA editing sites identified in the iMicroglia cohort (n=27). **B)** RNA editing rate by editing indexes and stimulation status. P-values from two-sided paired t-test. ns: non significant; ***: P < 0.001; ****: P < 0.0001. **C)** RNA Editing rate differential analysis comparing LPS-stimulated and basal samples. **D)** Comparing effect sizes of DESs identified under LPS stimulation within the discovery cohort and the replication cohort plotted.**E)** As in **(C)**, but comparing IFN*γ*-simulated and basal samples. 155 A-to-I and 17 C-to-U down-regulated and 2,330 A-to-I and 64 C-to-U up-regulated sites were found. **F)** As in **(D)**, but under IFN*γ* stimulation.

### RNA editing alters sequence stability both individually and combinatorially

A-to-I RNA editing is thought to largely reduce base pairing of complementary sequences, reducing structural stability. We then assessed whether the RNA editing sites responding to stimulation could alter structural stability using in *silico* secondary structure prediction. We reasoned that this may partly explain the observed associations between editing and differential expression. We clustered differentially edited sites by their nearby sites and made three types of comparison, comparing the predicted stability of a 2000 bp sequence window across a cluster of sites: 1) per-site editing, comparing a single base change at a specific site, considering each editing event separately; 2) per-gene editing, which aggregates all potential editing sites identified within a gene sequence; 3) combinatorial editing, whereas all possible combinations of editing events that can occur within a cluster are tested. Each combination represents a distinct pattern of edits that can be present simultaneously **(Figure 5A)**. RNA molecules tend to fold into structures with the lowest possible free energy, known as the minimum free energy (MFE), which can be predicted from sequence alone^37^. To assess the effect of RNA editing on sequence stability, we calculated per-sequence MFE for both unedited and edited sequences per gene. The difference between the two, referred to as ΔMFE, indicates the effect of editing, with a negative ΔMFE suggesting increased sequence stability post-editing. We observed that both IFNβ and LPS stimulation-induced sites affect the stability of the gene sequence in both directions **(Fig. 5B; Supplementary Table 14)**. We also confirmed that the ΔMFE increases positively in magnitude (more destabilizing) in proportion to the number of edited bases in a sequence, in line with expectation **(Fig. 5C; Supplementary Table 14)**. We highlight an example of a transcript with multiple editing sites, the *LIM* domain kinase 2 gene (*LIMK2*), which has five editing sites identified in the range chr22:31,276,967-31,277,313. The first two C-to-U sites and one A-to-I site are located within exons whereas the last two A-to-I editing sites were located within the 3’UTR **(Fig. 5D; Supplementary Table 15)**. The combination of five identified editing events and their corresponding ΔMFE shows that the accumulated effect of editing events is greater than that of a single editing event **(Fig. 5E; Supplementary Table 15)**. Interestingly, In *LIMK2*, the editing events combine to stabilize the secondary structure. The exonic C-to-U site at chr22:31,276,967, noted as position 1, has the single largest effect on RNA stability, with increasing sequence stability with the addition of sites at positions 3-5. Converting these four sites from the unedited to the edited bases *in silico* had a visible effect on the predicted secondary structure and overall confidence rate (**Figure 5F**). Another example for effects of RNA editing on gene structure is presented with the G protein-coupled receptor 141 gene *GRP141*, in which we observed six A-to-I sites (chr7:37,742,460-37,742,819), all located within the 3’UTR **(Supplementary Fig. 6A).** Unlike *LIMK2*, the editing events of *GPR141* act cumulatively to destabilize the structure. Interestingly, the 5th editing event (chr7:37,372,804: A-to-I) appears to drive the overall effect on the structure **(Supplementary Fig. 6B; Supplementary Table 31).** However, the secondary gene structure shows that while editing event 5 is changing the delta MFE significantly, its sole event does not have any considerable impact on the structure. **(Supplementary Figure 6C).** Thus, stimulation-responsive RNA editing can act to either stabilize or destabilize RNA structures.

**Figure 5.**
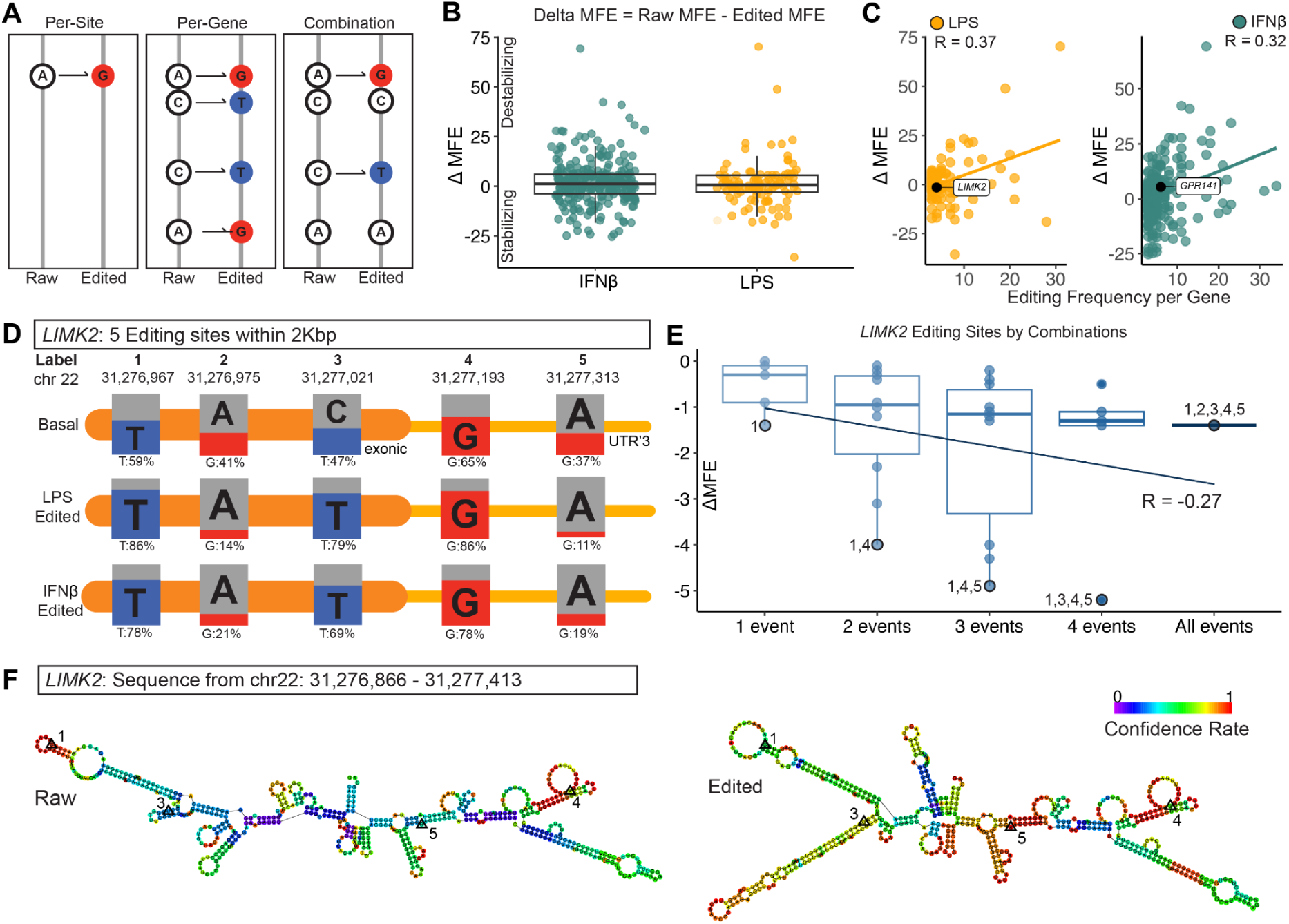
Predicted RNA stability changes by editing events in both simulations. **A)** Diagram of types of editing events modelled in silico. **B)** Delta Minimum Free Energy (ΔMFE) changes under stimulation. **C)** ΔMFE plotted against editing frequency per gene. **D)** Editing sites within *LIMK2* from chr22:31,276,967 to 31,277,313. The pair-editing percentage is noted for each basal, LPS-stimulated, and IFNβ-stimulated reads. **(E)** ΔMFE analysis for all possible combinations of editing events in *LIMK2*. **(F)** *LIMK2* gene structure diagram of unedited (raw) version and edited version with four events, labeled 1 (chr22:31,276,967), 3 (chr22:31,277,021), 4 (chr22:31,277,193), and 5 (chr22:31,277,313). Nucleotides colored by the calculated confidence of the predictability.

### Analysis of recoding events identifies an RNA mimic of a known genetic mutation

Across our three cohorts, we identified a total of 30,957 (29,466 A-to-I and 1,491 C-to-U) editing sites associated with 4,926 genes, of which 357 were annotated to be located within exons. Likely due to the aforementioned coverage differences, the discovery cohort showed the highest number of editing sites (165) in exons, and 17 sites were found in all three cohorts. Out of 357 exonic editing sites, 173 sites, associated with 107 genes, were differentially edited at least once under either stimulation. These differentially edited exonic sites were divided by editing index (74 A-to-I sites and 99 C-to-U sites) to observe their distribution of predicted codon substitution types **(Fig. 6A; Supplementary Table 16).** The majority of substitution types for these subset of exonic sites were detected to be nonsynonymous or synonymous SNV. However, two A-to-I sites, each associated to *SDHD* and *SLC30A4,* were identified with stoploss substitution, and two C-to-U sites identified with *HSP90B1* and *APOBEC3F* were associated with stopgain substitution **(Fig. 6B; Supplementary Table 17)**. We then studied the nonsynonymous recoding events of the differentially edited exonic sites by identifying their specific amino acid base before and after the editing. We categorized the recoding types into two groups: significant, where alterations in the resulting protein’s amino acid sequence impact its structure and function, potentially affecting biological processes and disease outcomes, and benign, where changes in the amino acid base do not significantly alter the original protein’s function. Among the exonic sites that were differentially edited, 47 sites were predicted to have significant amino acid recoding events, while amino acid base change of 121 sites were benign **(Fig. 6C; Supplementary Table 17)**. 12 sites with identified recoding events were previously reported in REDIportal, leaving 161 sites as novel recoding event discoveries. 11 sites, associated with 9 genes, were found to be differentially edited in all three cohorts, among which 3 sites were found with significant recoding events.

**Figure 6.**
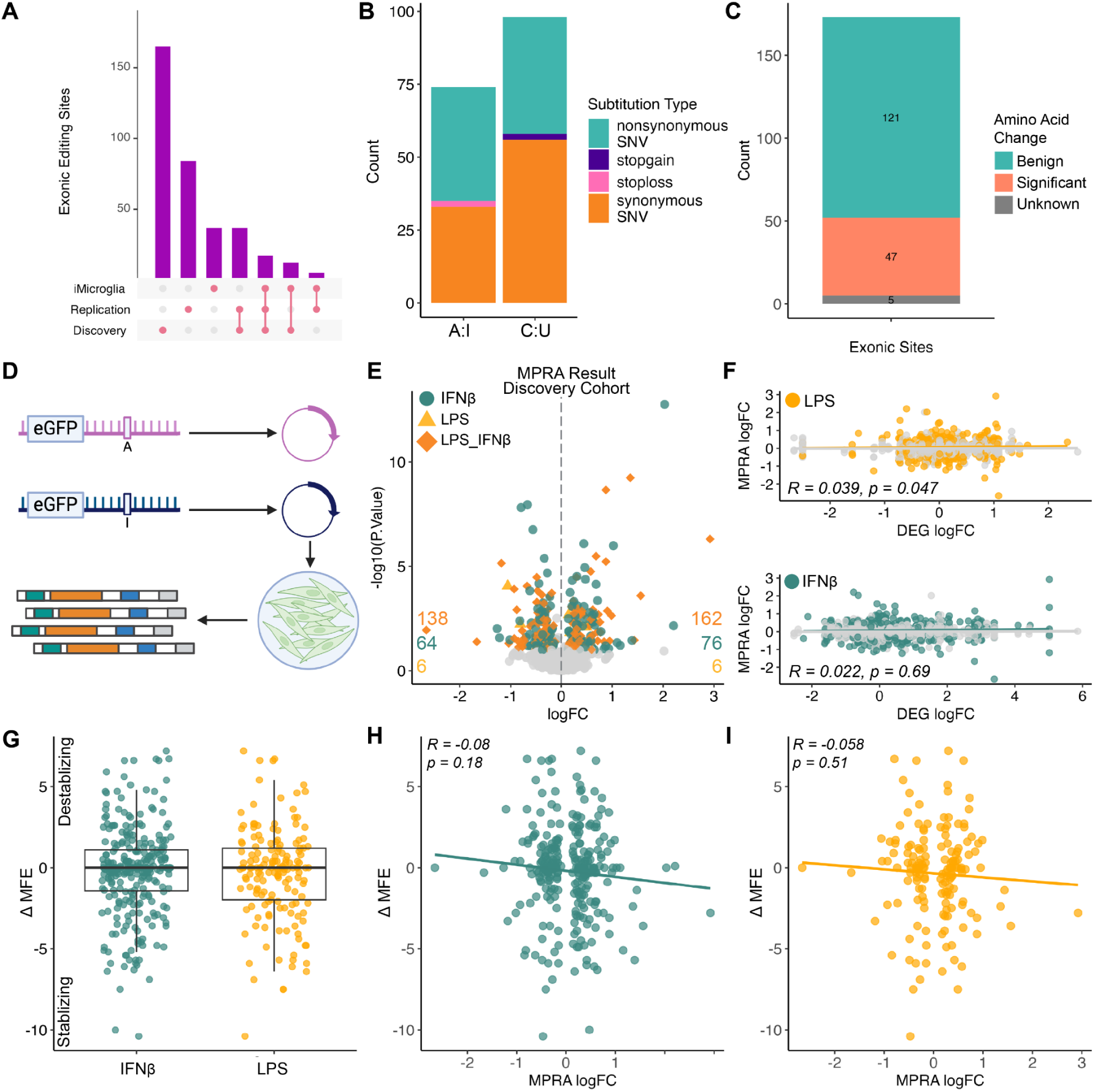
MPRA validation of 3’UTR sites. **A)** Counts of exonic editing sites identified in each subset of cohorts and overlapping between cohorts. **B)** Exonic editing sites that were differentially edited at least in once under either stimulation across all cohorts. **C)** Predicted amino acid recoding changes for the 173 exonic editing sites that were differentially edited at least in once under either stimulation across all cohorts. **D)** Experimental design of MPRA to identify functional 3’ UTR RNA editing sites that regulate mRNA abundance in HEK293T cells. **(E)** MPRA analysis result for editing sites identified in the Discovery cohort under IFNβ stimulation and LPS stimulation. **(F)** logFC of MPRA results against DEG analysis logFC under IFNβ stimulation for the identical sites, followed by sites identified in the discovery cohort under LPS stimulation. **(G)** Delta MFE of sequences 150 bp flanking the 290 editing sites (300 bp total), associated with 138 genes, identified as significant in MPRA analysis, under IFN stimulation, and for 162 sites associated with 84 genes under LPS stimulation. **(H)** Calculated ΔMFE against logFC of the same site from the MPRA analysis result. Each dot represents a 300bp sequence. **(I)** Same as (**H**) but for those sites identified under LPS stimulation.

We reasoned that some of the observed recoding events may alter protein structure or function, and therefore may mimic previously reported genetic missense mutations^38^. Our pipeline specifically excludes RNA editing sites where the alleles precisely overlap known common and rare variants, but we note that RNA editing of adjacent nucleotides in a codon could still act to cause a missense mutation. 107 genes associated with 168 differentially edited exonic sites at least once in three cohorts were detected with 121 benign recoding and 47 significant recoding events. We proceeded to look up whether any of the 47 recoding events have been reported as DNA mutations in the ClinVar mutation database. One C-to-U recoding event (chr5:78984968:C:T) identified in the discovery cohort in the Arylsulfatase B (*ARSB*) gene, p.Ser94Leu, was previously reported as a cause of the recessive lysosomal storage disorder Mucopolysaccharidosis type 6, which results from a deficiency of the Arylsulfatase B protein. The C-to-U editing of this site increases from a mean of 2.6% in baseline to 54% under IFNβ stimulation, effectively inducing a heterozygous loss-of-function mutation. These results suggest that recoding RNA editing events upregulated by stimulation can potentially act to mimic pathogenic mutations.

### In vitro validation of 3’UTR editing using a Massively Parallel Reporter Assay

Finally, we further investigated RNA editing sites observed within 3’UTRs, which can have specific effects on regulating mRNA stability, through altering the binding sites of RNA-binding proteins^39^ or microRNAs^17^, or altering double-stranded RNA formation^40^. Across three cohorts, we identified 2,150 sites (both A-to-I and C-to-U) located within 3’UTRs for testing using MapUTR^41^, a massively parallel reporter assay (MPRA). MapUTR screens for 3′ UTR variants affecting mRNA abundance post-transcriptionally, i.e. mRNA stability. We identified 533 sites with significant effects (FDR < 0.05) on 3’UTR mRNA abundance. For further downstream analysis, we focused on the sites identified in the discovery cohort under either stimulation. 300 sites were found to be significantly edited under both LPS and IFNβ stimulation, while 140 sites were significant under IFNβ stimulation and only 12 sites under LPS stimulation **(Fig. 6E; Supplementary Table 18)**. This suggests that sites found to be differentially edited under both stimulation are more likely to have significant effects on stability. When we compared the gene expression logFC of these significantly edited sites under LPS stimulation with the stability logFC results from the MPRA analysis, it had a negligible positive correlation (*R =* 0.039, *p =* 0.047) **(Fig. 6F; Supplementary Table 18)**.

We repeated the analysis for the sites found under IFNβ stimulation, identifying 290 significant sites out of 649 tested sites. Correlating the logFC of MPRA results and DEG analysis shows a non-significant correlation (*R =*0.022, *p =* 0.69) **(Fig. 6F; Supplementary Table 18).** Taken together, this suggests that differential expression is a poor predictor of changes in 3’UTR mRNA stability. We then took each site that was significant in MPRA analysis for sequence stability analysis, where we compared the MFE of the 300bp around 440 detected editing sites (301 bp input per site), using the sequence with and without the specific RNA editing event to mimic the MPRA *in silico* **(Fig. 6G; Supplementary Table 19)**.

For the sites identified under IFNβ stimulation, 283 unique sites were associated with 136 unique genes. We calculated ΔMFE of the 283 sites, extending the flanking sequence to 150bp before and after the targeted editing site, and also compared the ΔMFE to the logFC result of MPRA analysis for each corresponding site (*R* = -0.08, *p* = 0.18) **(Fig. 6H; Supplementary Table 20).** Then we repeated the analysis for 157 sites associated with 83 genes identified under LPS stimulation (*R* = -0.058, *p* = 0.51) **(Fig. 6I; Supplementary Table 21).** In summary, while we could identify a functional association between 3’UTR editing sites and RNA stability *in vitro*, we could not observe an explanation for these effects either through differential gene expression or *in silico* sequence stability prediction, suggesting an independent mechanism behind these associations.

## Discussion

In this study, we combined 297 RNA-seq samples from 3 cohorts to assess the effect of LPS and IFN stimulation on both A-to-I and C-to-I RNA editing in myeloid cells. Our large sample size has empowered us to assess correlations with differential gene expression and prioritize specific genes and editing sites for further study, while also revealing the technical complexities of accurately assessing RNA editing from RNA-seq data.

We identified widespread differential editing in response to inflammatory stimulation and observed that C-to-U sites as a group have a larger effect size than A-to-I sites across all three cohorts. Pro-inflammatory stimuli such as LPS and IFN induce widespread changes in gene expression affecting a large number of genes and pathways. We therefore expect RNA editing changes to represent a relatively small proportion of the total affected genes. In line with that expectation, we identified a small but significant overlap between genes affected by both differential expression and differential editing, with a largely concordant direction of effect between the two phenomena. Our results suggest a shared mechanism between editing and expression, although we are unable to identify a precise causal relationship.

One of the strengths of our study is the use of three independent cohorts of myeloid cells, generated using different RNA-seq library preparations. This allows us both to replicate our findings and to highlight the innate challenges in identifying RNA editing from RNA-seq data. While replication of identified sites was high, differences in library preparation and RNA quality between our discovery and replication cohorts, and even within different batches of the same cohort, meant that C-to-U sites were less likely to be replicated than A-to-I sites. Unlike A-to-I sites, which are maintained in the REDiportal database, there is currently no corresponding resource for C-to-U. Our overlap with a previously published study^20^ was small, and this could be due to differences in sample size, technical differences in sequencing, or technical difficulties with retrieving the correct coordinates. We suggest that a future C-to-U-focussed database pay close attention to library preparation parameters due to these clear technical biases.

The question of the functional impact of RNA editing is an important one for the field. Here we used both *in silico* predictions of structural stability and *in vitro* modelling of 3’UTR RNA abundance to assess the capacity of both types of stimulation-responsive editing to alter RNA stability. While both methods predict that specific sites would affect stability, the two predictions largely disagree and neither can be well explained by the observed differential expression of the host mRNA. A further potential outcome of RNA editing is to recode the proteome. Using our set of putative recoding events we identified a site that mimics a known mutation in *ARSB*. Future investigation into recoding sites could identify additional mutation mimics or the creation of neoantigens^42^.

A limitation of our study is the lack of a disease comparison due to a lack of power. Aberrant RNA editing has been observed in multiple diseases with a potential myeloid component ^43–45^ Furthermore, we did not have corresponding genetic data to assess the genetic regulation of RNA editing, so-called editing quantitative trait loci (edQTLs). We note that while several studies have mapped edQTLs in patient tissues^43–45^, these studies have neglected C-to-U editing. Future work could apply an edQTL framework to both types of RNA editing in myeloid cells to identify new causative genetic effects on disease.

## Methods

### Sample Collection

This study is part of the MyND (Myeloid cells in Neurodegenerative Diseases) initiative, which aims to characterize the immune system (monocytes and microglia) from Parkinson’s disease, Alzheimer’s disease, and age-matched controls^1^. We collect fresh blood samples from various cohorts and institutions across New York City, including Mount Sinai Movement Disorders Clinic (MSMD), NYU Parkinson’s and Movement Disorders Center (NYUMD), Beth Israel Movement Disorders (BIMD), Center for Cognitive Health (CCH), and Alzheimer’s Disease Research Center (ADRC). Across discovery and replication cohorts, a total of 90 unique donor samples were acquired.

### Discovery Cohort

The discovery cohort comprises 55 donors, categorized into 11 with Alzheimer’s Disease (AD), 11 with Parkinson’s Disease (PD), 11 with Mild Cognitive Impairment (MCI), 11 Young Controls (YC), and 11 Controls. The cohort consisted of 33 female and 22 male donors, with ages ranging from 23 to 85, with median age of 72. Blood was diluted with twofold PBS (Gibco) and subjected to the isolation of Peripheral Blood Mononuclear Cells (PBMCs) using SepMate tubes (StemCell Technologies) filled with 15 ml of Ficoll-Paque PLUS (GE Healthcare) through 15-minute centrifugation at 1,200g. The PBMCs were then washed with PBS and cryopreserved in liquid nitrogen. Cryopreserved PBMCs were thawed in 10 ml of media (RPMI + 10% FBS + P/S + Gln) and monocytes were isolated using the CD14+ selection kit, which uses CD14+ magnetic beads (Miltenyi) to isolate CD14+ monocytes characterized by the expression of the CD14 cell surface receptor. PBMC and monocyte viability was assessed using a CountessII Automated Cell Counter (Thermo Fisher) (Navarro et al. 2021). 10,000 monocytes/well were cultured at a concentration of 1 million cells/ml using RPMI media supplemented with antibiotics, Glutamine (Gln) and Fetal Bovine Serum (FBS). The treatments, LPS 10 ng/ml and IFNβ 20 ng/ml were added to the wells separately. For untreated samples, the same volume of PBS was added. Samples were cultured for 24 hrs at 37 C, washed in PBS, centrifuged, and resuspended in 350 µl of RLT + 1% 2-Mercaptoethanol (Sigma Aldrich) and stored at -80 °C for downstream RNA isolation.

### Replication Cohort

The replication cohort consisted of 35 donors. 20 donors had a diagnosis of PD and 15 were from controls. The cohort included 16 female and 19 male donors, with ages ranging from 45 to 79 and a median age of 68. Experiments were performed in 2 independent experimental batches. PBMC isolation was performed following the same protocol as described for the Discovery Cohort. Monocytes were isolated from cryopreserved PBMCs as follows. Cryopreserved PBMCs were thawed in a water-bath and diluted in 10 ml of media (RPMI + 10% FBS + P/S + Gln) and centrifuged . After washing, PBMCs were sorted to monocytes using CD14^+^ magnetic beads (Miltenyi) in the Manual sorter following the manufacturer’s instructions. Monocyte number and viability was assessed using Countess II Automated Cell counter (Thermo Fisher). After sorting, monocytes were resuspended in supplemented RPMI media at a concentration of 1 million cells/ml. 500,000 cells were plated with the following stimuli: LPS-EK ultrapure (InvivoGen) 10 ng/ml, IFNγ (R&D Systems) 20 ng/ml or baseline (media + PBS). After 24 hrs of culture at 37 °C and 5% CO2, cells were washed with PBS and pellets were resuspended in 350 µl of RLT + 1% 2-Mercaptoethanol (Sigma Aldrich) and stored at -80 °C for downstream RNA isolation.

### iMicroglia Cohort

Isogenic iPSCs for G2019S (wild-type, heterozygous, and homozygous clones from one donor) were provided by the Cookson Lab^46^. Genotype was validated by Sanger sequencing, and all clones displayed a normal karyotype and standard pluripotent expression markers. iPSCs were maintained in StemFlex medium and cultured on Matrigel-coated 6-well plates. Media was changed every other day, and cells were split using ReLeSR about once per week^29^.

iPSCs were first differentiated to hematopoietic stem cells over a period of 12 days. Briefly, iPSCs were dissociated into a single-cell suspension using TrypLE Select and plated at ∼500k cells/well onto tissue culture-treated 6-well plates in StemFlex + rock inhibitor. After 24 hours (HPC diff: day 1), the media and floating cells were collected, spun, and resuspended in differentiation medium (DM) 1, which consisted of HPC base medium supplemented with FGF2, BMP4, Activin A, LiCl, and rock inhibitor, and replated and cultured under hypoxic (5% O2) conditions. On day 3, cells were switched to DM 2, which included FGF2 and VEGF. On day 5, cells were switched to DM 3, which included FGF2, VEGF, IL6, TPO, SCF, and IL3, and switched back to a normoxic incubator. Cells were supplemented with additional DM3 every other day until floating HPCs were collected between days 11-15 and either frozen in Bambanker Freezing Medium or cultured directly in iMGL medium as described below.

HPCs were plated at 166k/well in a 6-well plate in iMGL medium, consisting of DMEM/F12, GlutaMAX, Non-essential amino acids, B27, N2, ITS, insulin, and MTG, and supplemented with fresh IL-34, M-CSF, and TGFb on each feeding day. Cells were fed every other day with 1mL medium, and were split 1:2 once per week over the course of the 25 day HPC → iMGL differentiation protocol. At day 25-28, cells were supplemented with the above factors along with two maturation factors: CX3CL1 and CD200. Cells were cultured in this medium for 3 days and used for subsequent assays on day 28-31. For RNA-seq and qPCR analysis: matured iMGLs were plated at 500k cells per well of a 6-well plate and treated with 10ng/mL of LPS or 20ng/mL of IFNγ. After 24 hours of treatment, cells were washed once with PBS and then lysed directly in the well using RLT buffer with beta-mercaptoethanol (Qiagen RNA-easy Mini Kit) and lysed cells were kept at -80 C and processed together. Each staggered HPC → iMGL differentiation is considered one biological replicate^29^.

### RNA sequencing

RNA was isolated using RNeasy Mini kit (Qiagen) following the manufacturer’s instructions and including the optional DNase treatment. RNA was stored at -80 °C prior to library preparation. Library preparation and sequencing was performed at Genewiz Inc. using the SMART-Seq v4 Ultra Low input library preparation protocol, which uses poly-A selection. The samples in discovery and iMicroglia cohorts were sequenced with an average depth of 30 million 150-bp paired-end reads using the Illumina HiSeq 4000 platform. The replication cohort was sequenced to a depth of 66 million mapped reads with a median read length of 115 bp. We processed the FASTQ files with RAPiD, an RNA-seq processing pipeline implemented in the Nextflow framework. The pipeline aligns the reads to the hg38 genome built with STAR (v.2.7.2a)^47^ using the GENCODE v.30 transcriptome reference. Gene expression was quantified with RSEM (v1.3.1)^48^. Pre-alignment sequencing quality and technical metrics were assessed using FASTQC (v0.11.8). Post-alignment quality control using Picard (v2.22.3)^32^ and Samtools^49^ provided essential metrics for evaluating the quality of sequenced data. Initial inclusion criteria of post-alignment quality control consisted of at least 20 million passed reads, with at least 20% of reads mapping to coding regions and ribosomal rate < 30%. Four samples were removed based on sex mismatches. Outliers were also removed after adjusting for covariates using dimensionality reduction through principal component analysis (PCA) that were selected to be used in differential analyses. Seven samples were removed for being PCA outliers.

### RNA editing analysis

We constructed an RNA editing pipeline to process the input BAM and metadata files through a series of steps to output an annotated site-specific RNA editing rate matrix associated with each sample. The pipeline employs two major computational tools: JAVA framework for accurate SNV assessment (JACUSA2)^32,50^ and ANNOVAR^51^. JACUSA extracts RNA editing sites and their read coverage from RNA-seq data and applies a statistical model to estimate the probability of RNA editing at each site. We ran JACUSA (v.2.0.1) to find both *de novo* sites and a set of known sites from REDiPortal (v2.0)^32^. *De novo* sites were filtered with >10 read coverage and > 3 coverage of edited allele, whereas known sites were filtered with >5 read coverage and >3 coverage of edited alleles. After concatenating all sites and samples into a single matrix, we applied group-wise filters. For the discovery and replication cohort, the filters were set so that each site must be detected in at least 10 donors, with a minimum editing rate of 0.01 with 10% site and sample missingness. For the iMicroglia cohort, these filters were lowered to 2 samples, while keeping the editing rate and site/sample missingness threshold the same. Sites were then annotated with overlapping common variants, and segmental duplications were filtered out. Finally, the final set of consensus sites are re-called in each sample using JACUSA, to resolve sites with true missingness (0 reads in that sample) from samples that failed the individual read coverage thresholds on the first pass. Overall, the pipeline outputs three separate matrices: a sample-level coverage matrix, sample-level editing matrix where RNA editing levels for each sample are provided as a proportion - the count of edited reads divided by the total reads at that specific nucleotide position, and site-level annotation matrix. We calculated an editing discovery rate for A-to-I and C-to-U sites by counting all non-zero editing sites of that type in a sample divided by the total mapped bases in millions.

### Differential editing analysis

We split out A:G/T:C (A-to-I sites on either strand) from C:T/G:A (C-to-U sites) variants to examine sources of technical and biological variation on the two editing types. We first calculated the first 10 principal components and performed univariate correlations of each principal component with known clinical variables, the expression of *ADAR* and *APOBEC* genes, and sequencing quality metrics derived from Picard. We then applied VariancePartition (v1.28.7)^52^, which uses linear mixed modeling to jointly assess the contribution of each variable to the variance in each editing site. We then conducted a linear model-based differential expression analysis between the stimulated and basal samples, using the limma (v.3.60.6)^53^. All editing ratio values in the metadata were log-normalized, and any missing values (corresponding to no detectable sequencing reads at that position in that sample) were set to zero. Any editing site with a standard deviation of zero across the samples was removed. For the discovery cohort, two models were used for differential analysis: one with only selected categorical covariates and another including those same covariates along with *ADAR* and *APOBEC* expression in TPM. The second model aimed to explore whether alterations in RNA editing rates between groups was attenuated by accounting for the variable expression in editing enzymes. The contrasts were made on the condition (IFNβ, LPS) vs. basal. For the discovery cohort, model 1 was *∼ stimulation status + diagnosis + age + sex + ribosomal base coverage percentage*, and model 2 was model 1 equation + TPM values of *APOBEC3G, APOBEC3F,* and *APOBEC3A,* identified through principal component analysis (PCA) and variancePartition analysis. Model 1 of the replication cohort was kept the same as that of the discovery cohort with addition of variable *batch*. The iMicroglia cohort were modelled using *∼ stimulation status + genotype*.

For comparing differential analysis in RNA editing rate across all cohorts, it was expected that the effect size measured by log_2_ fold change (logFC) values of these sites would be small. Therefore, no logFC threshold was applied when determining a site’s statistical significance. A site with an adjusted P value < 0.05 was considered a significantly differently edited site (DES).

### Differential gene expression

To ensure the quality of the gene expression differential analysis, we initially filtered out genes with mean TPM values lower than one across all samples in the discovery cohort. In the discovery cohort, 58,929 genes were initially quantified, of which 12,961 genes passed the TPM > 1 criterion. Then, differential expression analysis was done with R packages edgeR (v 4.2.2) ^54^ and Voom (3.60.6) ^55^. The model used for analyzing samples in the discovery cohort was *∼ stimulation status + batch + sex + age*. Statistically differentially expressed genes (DEGs) were identified based on a threshold of adjusted P < 0.05 and logFC > 1.

### *In silico* stability analysis

To assess the effect of RNA editing on sequence stability, we calculated per-sequence Minimum Free Energy (MFE), a thermodynamic measure of RNA stability^37^ . We calculated MFE both for unedited-sequence and per-gene edited sequence using RNAfold (v 2.4.12) ^56^ from the Vienna RNA Package^56^. ΔMFE was calculated by getting the difference between MFE of raw and edited sequence, with a negative ΔMFE implying that editing increases stability, and *vice versa*. We visualized the predicted structures with and without edited nucleotides using the RNAfold web server.

### Massively Parallel Reporter Assay

We assessed whether the differentially edited sites found within 3’UTRs had an effect on RNA abundance *in vitro* using a massively parallel reporter assay (MPRA) design which uses a library of 164bp oligomers cloned into a reporter vector with either the reference or edited nucleotide. We aimed to test whether these identified 3’UTR editing events impact mRNA abundance using MapUTR^41,57^. Reporter plasmids carrying the same 3’UTR sequence, differing only at specific editing sites, are electroporated into HEK293T cells. After 24 hours, the mRNA was extracted and used to generate RNA-seq libraries. DNA-seq libraries were also generated from the plasmid library to allow for RNA normalization. Three biological replicates were collected. Following sequencing, we used MPRAnalyze^58^ to identify sites with differential RNA/DNA ratio between the edited and unedited alleles. FDR ≤ 0.1 and |logFC| ≥ 0.1 were used to call significance.

### Recoding event analysis

For all editing sites identified in exonic regions across the three cohorts, we used ANNOVAR to annotate the amino acid base changes. If the identified editing site was neither a common nor a rare SNP by comparing the site with the *hg38_dbsnp153Common* database, we determined the amino acid at that location before and after the editing event. Amino acids were categorized as hydrophilic (A, G, I, L, P, V, M), hydrophobic (F, W, Y), basic (R, H, K), acidic (D, E), and polar uncharged (S, T, N, Q, C). If the editing event caused a change in an amino acid group, it was considered a significant change in the sequence. If the change did not alter the amino acid group, it was labeled benign. Additionally, if the amino acid base change due to the editing event had been previously reported (found in the REDIportal database v2.0), it was considered a known event. If there was no match in the REDIportal database, the event and the resulting amino acid change were considered novel findings.

## Data Availability

RNA-seq samples for the discovery and replication cohorts of stimulated microglia have been deposited on dbGAP (accession ID: phs002400.v1.p1) and will be released upon publication. The iMGL cohort RNA-seq samples have been deposited on the Gene Expression Omnibus (accession: GSE240907).

## Code Availability

RNA editing pipeline: https://github.com/RajLabMSSM/editing-pipeline/tree/master/jacusa-pipeline Code for generating all figures: https://github.com/RajLabMSSM/RNA_Editing_Monocytes/

### URLs

RNAfold server: http://rna.tbi.univie.ac.at/cgi-bin/RNAWebSuite/RNAfold.cgi

## Acknowledgements

We thank the patients and families who donated material for these studies. This study was supported by the following National Institutes of Health grants: NIA U01-AG058635, NIA R21-AG063130, NIA R01-AG054005, NIA U01-AG068880, NIA RF1-AG065926, NIA R56-AG055824, NIA P30-AG066514, NINDS U54-NS123743, and NINDS R01-NS116006 to TR, JH, and HS. This work was supported in part through the computational resources and staff expertise provided by Scientific Computing at the Icahn School of Medicine at Mount Sinai and supported by the Clinical and Translational Science Awards (CTSA) grant UL1TR004419 from the National Center for Advancing Translational Sciences. Research reported in this paper was supported by the Office of Research Infrastructure of the National Institutes of Health under award number S10OD026880 and S10OD030463. EN is supported by the Spanish Ministry of Science and Innovation (PID2022-139936OA-I00). The content is solely the responsibility of the authors and does not necessarily represent the official views of the National Institutes of Health.

## Author contributions

JH and TR conceived and designed the study. HS analyzed the data, performed the statistical analyses, and created all figures, with technical assistance from WHC, EN, TF, and MSB. HS and JH wrote the manuscript. All sequencing data was generated by EN, AGE, AA, and MP. The MPRA experiment was performed by TF and GX. All authors have read and approved the final manuscript. JH and TR supervised the project.

## Supplementary Figures

**Supplementary Figure 1.**
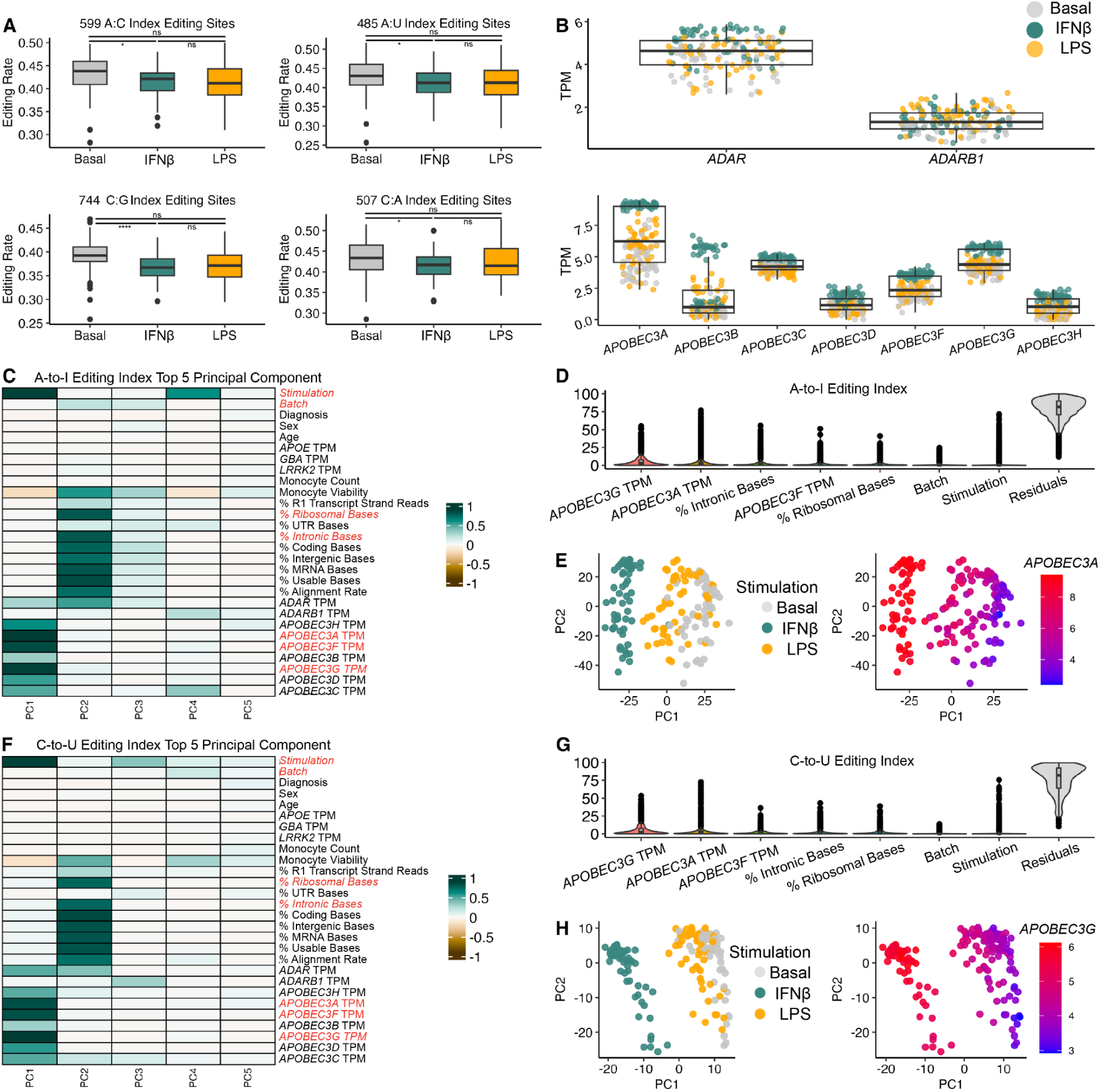
Editing rate variance in discovery cohort influenced by technical, biological, and experimental variables. **A)** Mean editing mean rates for 599 A-to-C, 485 A-to-U, 744 C-to-G, and 507 C-to-A substitutions by stimulation status. P-values from paired t-test. ns: non significant; **: P < 0.01; ***: P < 0.001. **B)** Normalized TPM of *ADAR* and *ADAB1* genes by sample stimulation status. Normalized TPM of *APOBEC3 (A,B,C,D,F,G,H)* genes by sample stimulation status. **C)** Principal Component Analysis (PCA) revealed that stimulation status, percentage reads in ribosomal and intronic bases significantly influenced A-to-I editing rates. Within the *ADAR* and *APOBEC* gene families, *APOBEC3A, APOBEC3F*, and *APOBEC3G* displayed the highest influence. **D)** Variance partition analysis on editing rate variance of A-to-I editing sites for covariates identified to be influential on editing rate. **E)** Principal component plot showing distinctions in samples by stimulation and by TPM of *APOBEC3A*. **F)** As for **(C)** but for C-to-U editing sites. **G)** Variance partition analysis on editing rate variance of C-to-U editing sites. **(H)** Principal component plot showing distinctions in samples by stimulation and by TPM of *APOBEC3G*.

**Supplementary Figure 2.**
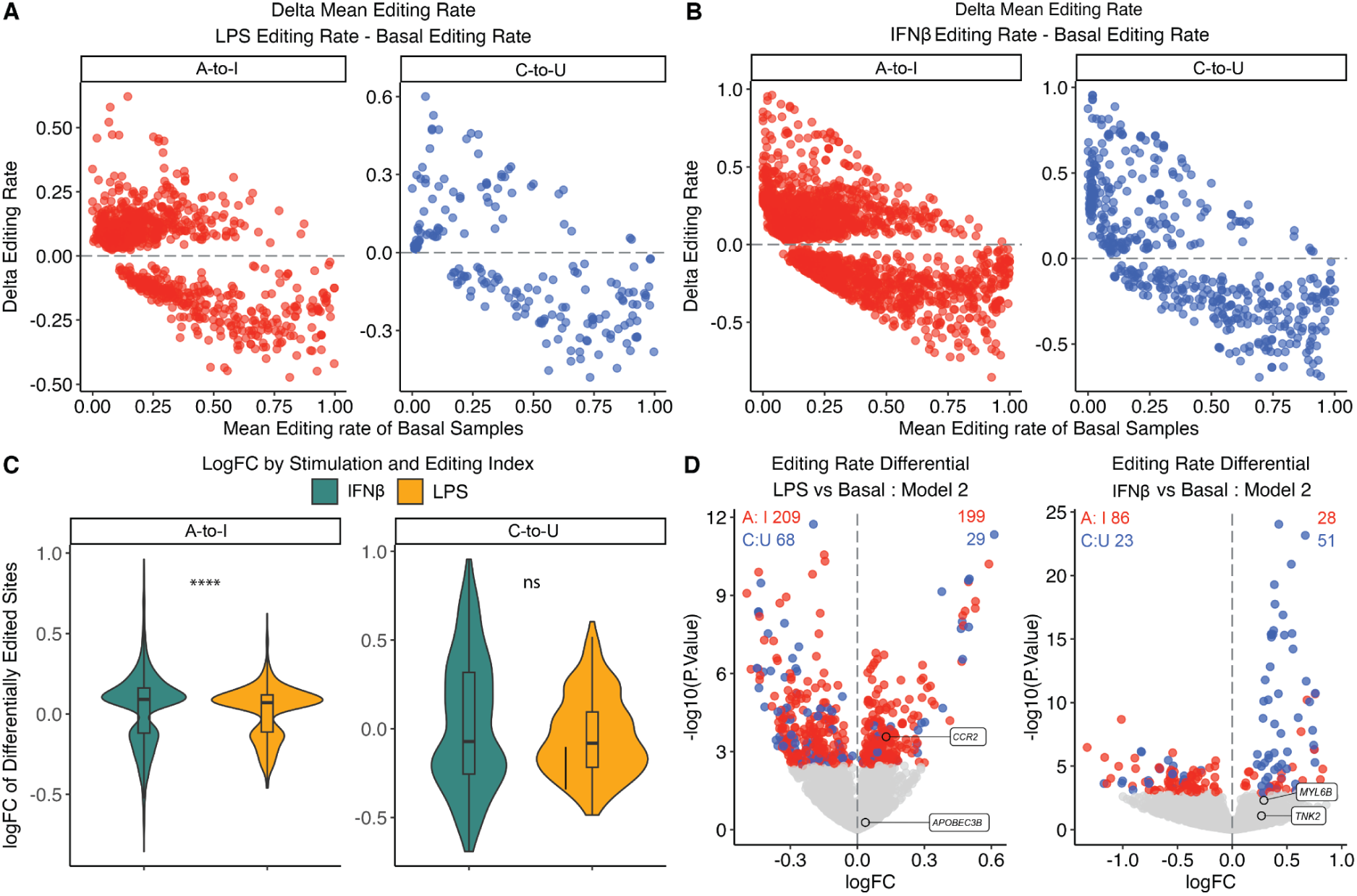
C-to-U editing sites have larger effect sizes in response to stimulation. **A)** Delta mean editing rate of differentially edited sites under LPS stimulation in the discovery cohort, plotted by editing indexes. (x axis: Mean editing rate of basal samples; y axis: delta mean editing rate (mean editing rate of stimulated sample - mean editing rate of basal sample at a site). **B)** As in (A) but under IFNβ stimulation. **C)** The effect sizes of 4,011 A-to-I and 670 C-to-U differentially edited sites were analyzed in the discovery cohort under IFNβ and LPS stimulation. For A-to-I editing events, the logFC differences between IFNβ and LPS were highly significant (****, P < 0.0001, under Wilcoxon test), while no significant difference was observed for C-to-U events. **D)** Differential editing analysis comparing simulated samples and basal samples when including *ADAR* and *APOBEC* gene expression as covariates. Under LPS-stimulation, 209 A-to-I and 68 C-to-U down-regulated sites and 199 A-to-I and 29 C-to-U up-regulated sites were found. Under IFNβ-stimulation, 86 A-to-I and 23 C-to-U down-regulated sites and 28 A-to-I and 51 C-to-U up-regulated sites were found.

**Supplementary Figure 3.**
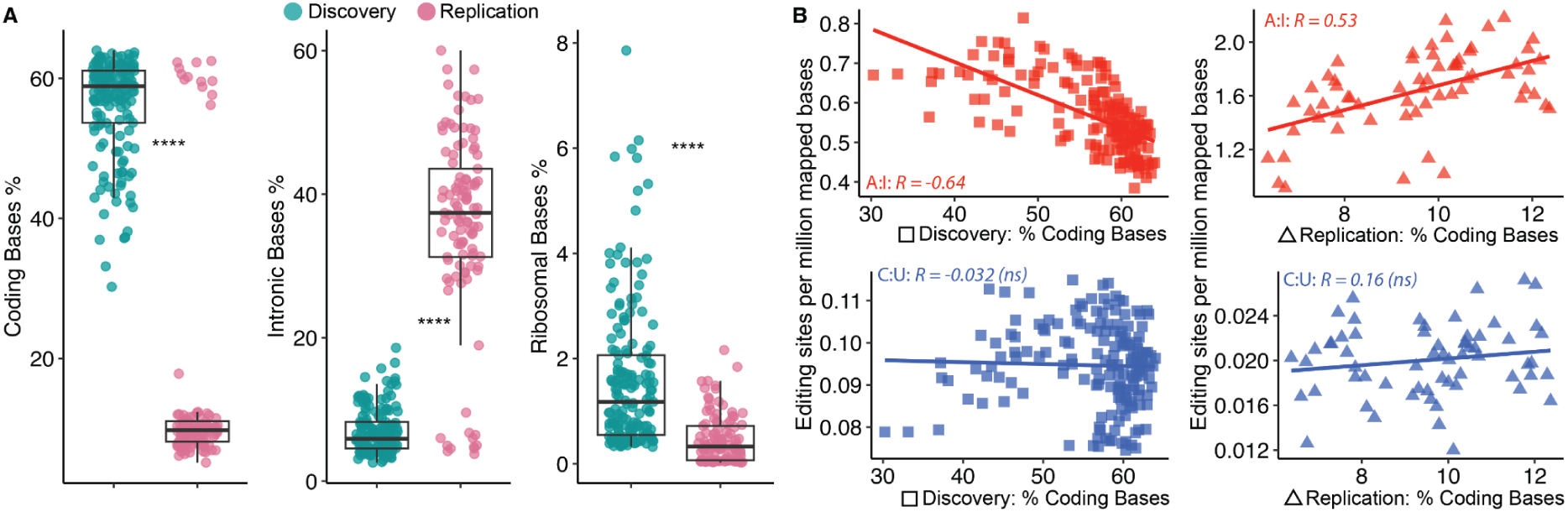
Library preparation differences explain differences in editing site discovery. **A)** Comparing % coding bases, % intronic bases, % ribosomal bases coverage of each sample in the discovery cohort and replication cohort. Discovery cohort: mean coverage of coding bases 56.27%, intronic bases 6.88%, and ribosomal bases 1.59%. Replication cohort: mean coverage coverage of coding bases 15.18%, intronic bases 35.69%, and ribosomal bases 0.47%. All comparison reports showed highly significant statistical results (****, P < 0.0001, under Wilcoxon test). **B)** Editing site discovery rate by percentage of coding bases coverage for each read for discovery (left) and replication (right) cohorts. In each unit of the plot, the Pearson correlation coefficient (r) and the associated p-value are printed.

**Supplementary Figure 4.**
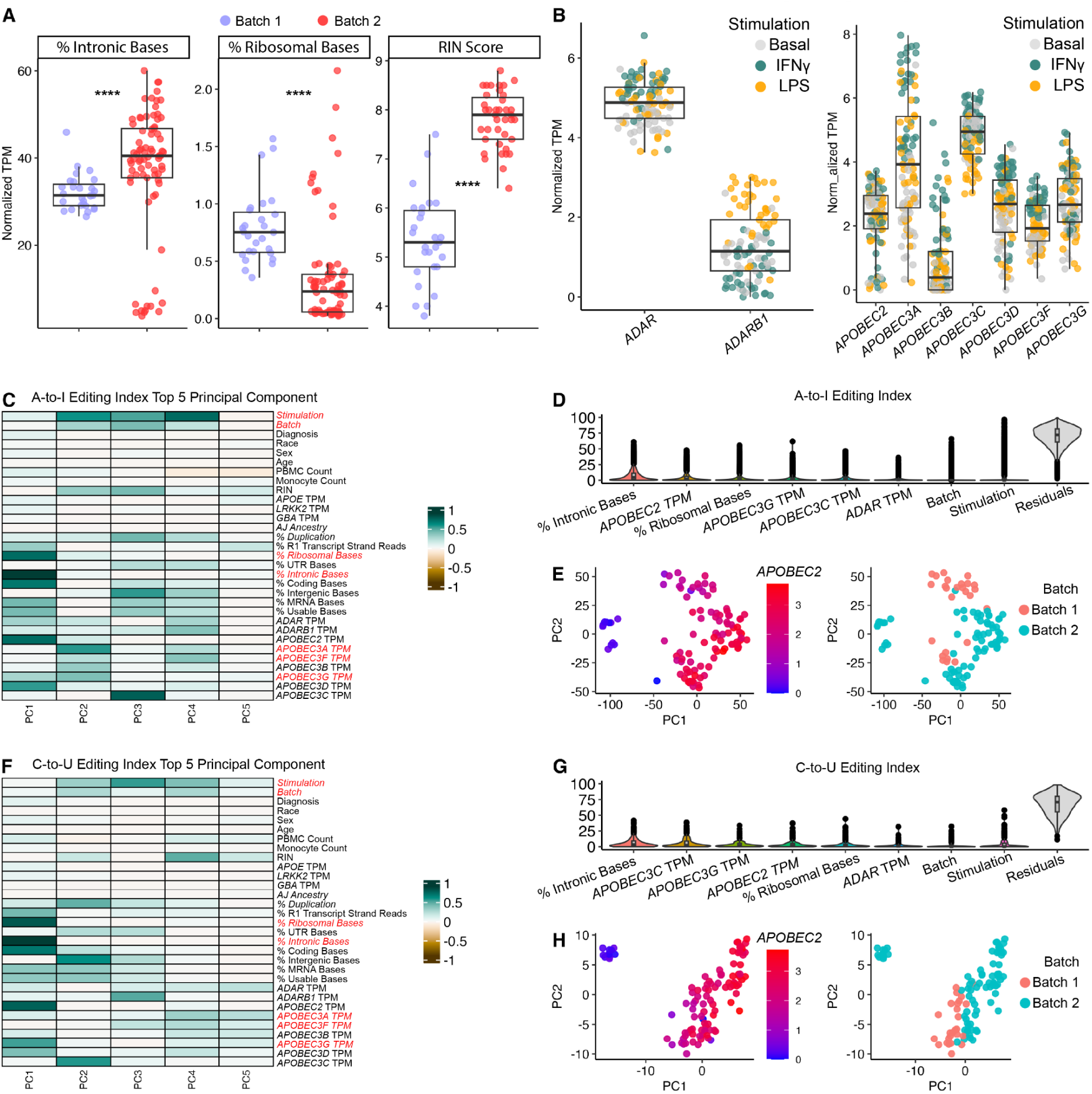
RNA editing in replication cohort influenced by stimulation status and *APOBEC* expression. **A)** A significant difference was observed in the percentage coverage of intronic bases, ribosomal bases, and RIN scores between batch 1 and batch 2 in the replication cohort. Statistical comparison results showed highly significant differences for intronic bases, ribosomal bases, and RIN scores (****, P < 0.0001, under Wilcoxon test) **B)** Normalized TPM of *ADAR, ADARB1, APOBEC2,* and *APOBEC3 (A,B,C,D,F,G)* genes by sample stimulation status. **C)** PCA of A-to-I editing rate variance, showing importance of stimulation status, batch, percentage reads in ribosomal and intronic bases. Within the *APOBEC* gene families, *APOBEC2, APOBEC3C*, and *APOBEC3G* displayed the highest influence. **D)** Variance partition analysis on editing rate variance of A-to-I editing sites. **E)** Principal component plot showing distinctions in samples by TPM of *APOBEC2* and batch. **F)** As of (C) but for C-to-U editing sites. **G)** Variance partition analysis on editing rate variance of C-to-U editing sites. **H)** As of **(C)** but for C-to-U editing sites.

**Supplementary Figure 5.**
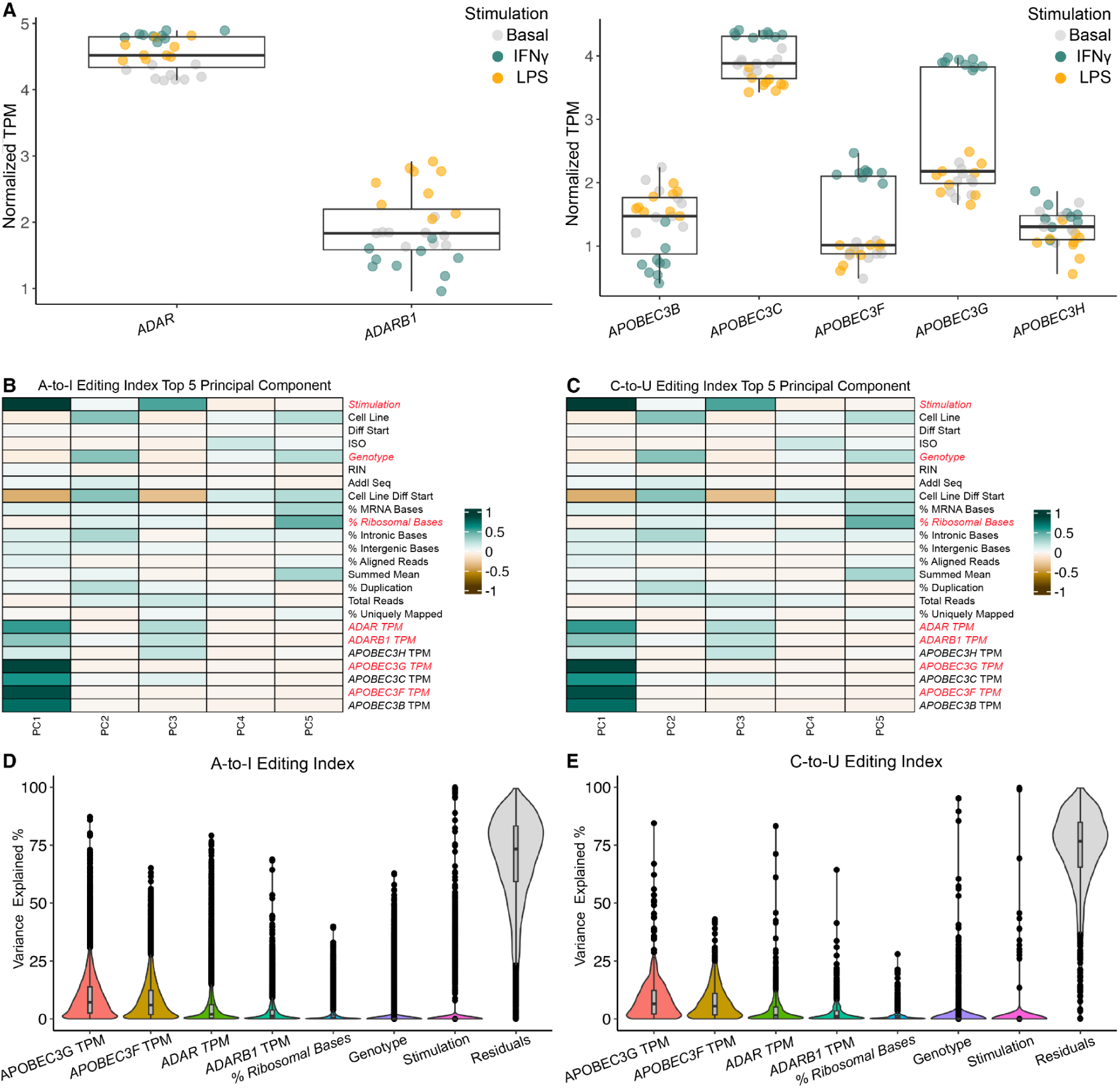
Editing analysis in the iMicroglia cohort. **A)** Normalized TPM of *ADAR, ADARB1, APOBEC2,* and *APOBEC3 (B,C,F,G,H)* genes by sample stimulation status. **B)** PCA revealed that stimulation status, percentage reads in ribosomal significantly influenced A-to-I editing rates. Within the *ADAR* and *APOBEC* gene families, *ADAR, ADARB1, APOBEC3G,* and *APOBEC3F* displayed the highest influence. Among technical covariatest, percentage coverage of ribosomal bases and sample stimulation status showed the highest influence on editing rate. **C)** As of **(B)** but for C-to-U editing rates. **D)** Variance partition analysis on editing rate variance of A-to-I editing sites. **E)** As in **(D)** but for C-to-U editing rates.

**Supplementary Figure 6.**
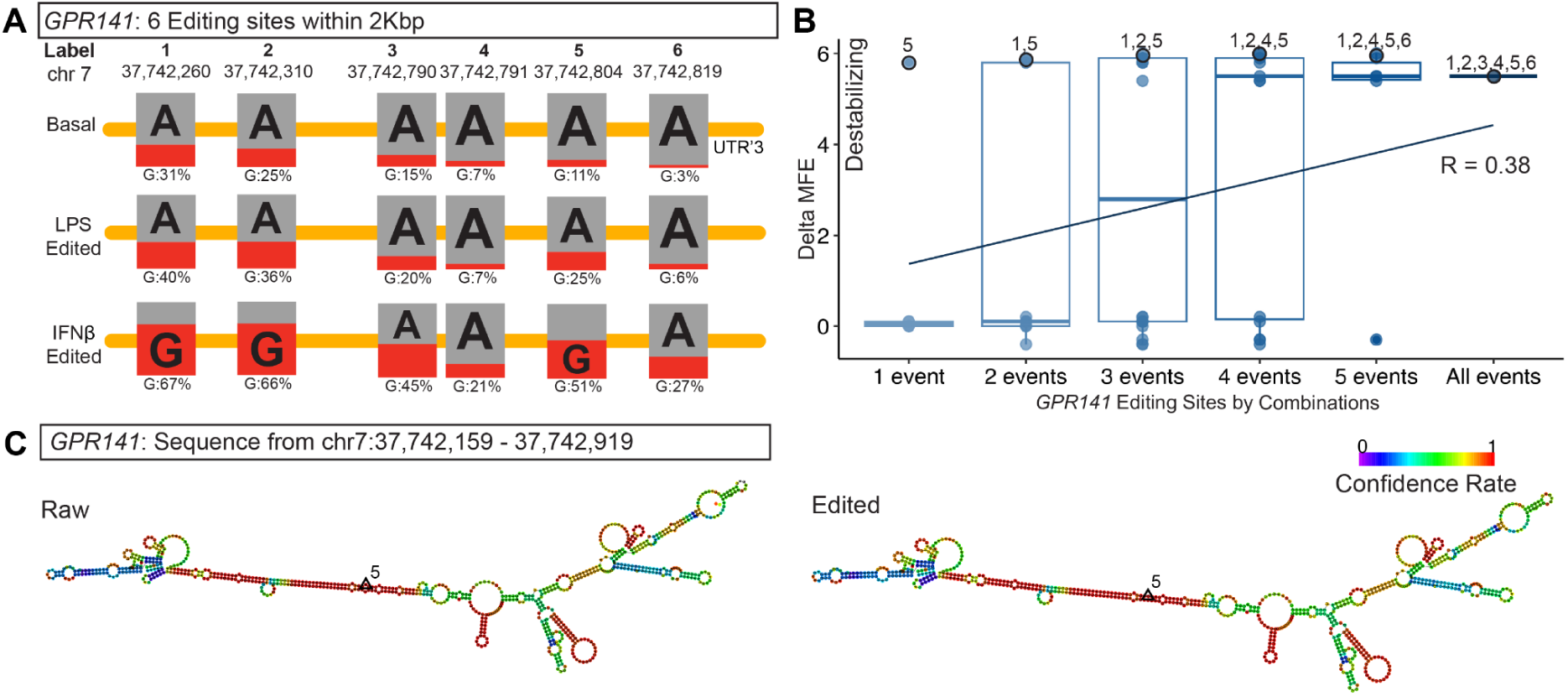
RNA sequence stability in *GPR141*. **A)** 3’UTR of *GPR141* from chr7:37,742,159 to 37,742,919. The pair-editing percentage is noted for each basal, LPS-stimulated, and IFNb-stimulated reads. *GPR141* was identified with six A-to-I editing events in these designated reads, and each event is labeled in number. **B)** Delta MFE analysis for all possible combinations of editing events in gene *GPR141*. Delta MFE for all possible combinations of six editing events were calculated individually and plotted. **C)** *GPR141* gene structure diagram of unedited (raw) version and edited version with the strongest editing event. The edited version of the gene structure reflects an editing site labeled 5, located in chr7:37,742,804.

## References

1. Navarro, E. et al. Dysregulation of mitochondrial and proteolysosomal genes in Parkinson’s disease myeloid cells. Nat Aging 1, 850–863 (2021).

2. Kosoy, R. et al. Genetics of the human microglia regulome refines Alzheimer’s disease risk loci. Nat. Genet. 54, 1145–1154 (2022).

3. Young, A. M. H. et al. A map of transcriptional heterogeneity and regulatory variation in human microglia. Nat. Genet. 53, 861–868 (2021).

4. Humphrey, J. et al. Long-read RNA-seq atlas of novel microglia isoforms elucidates disease-associated genetic regulation of splicing. medRxiv (2023) doi:10.1101/2023.12.01.23299073.

5. Lopes, K. de P., et al. Atlas of genetic effects in human microglia transcriptome across brain regions, aging and disease pathologies. Preprint at 10.1101/2020.10.27.356113.

6. Lopes, K. de P., et al. Genetic analysis of the human microglial transcriptome across brain regions, aging and disease pathologies. Nat. Genet. 54, 4–17 (2022).

7. Ramaswami, G. et al. Accurate identification of human Alu and non-Alu RNA editing sites. Nat. Methods 9, 579–581 (2012).

8. Porath, H. T., Knisbacher, B. A., Eisenberg, E. & Levanon, E. Y. Massive A-to-I RNA editing is common across the Metazoa and correlates with dsRNA abundance. Genome Biol. 18, 185 (2017).

9. Slotkin, W. & Nishikura, K. Adenosine-to-inosine RNA editing and human disease. Genome Med. 5, 105 (2013).

10. Wang, C., Zou, J., Ma, X., Wang, E. & Peng, G. Mechanisms and implications of ADAR-mediated RNA editing in cancer. Cancer Lett. 411, 27–34 (2017).

11. Samuel, C. E. Adenosine Deaminases Acting on RNA (ADARs) and A-to-I Editing. (Springer Science & Business Media, 2011).

12. Patterson, J. B. & Samuel, C. E. Expression and regulation by interferon of a double-stranded-RNA-specific adenosine deaminase from human cells: evidence for two forms of the deaminase. Mol. Cell. Biol. 15, 5376–5388 (1995).

13. Behm, M. & Öhman, M. RNA editing: A contributor to neuronal dynamics in the mammalian brain. Trends Genet. 32, 165–175 (2016).

14. Bazak, L. et al. A-to-I RNA editing occurs at over a hundred million genomic sites, located in a majority of human genes. Genome Res. 24, 365–376 (2014).

15. Zhang, P. et al. On the origin and evolution of RNA editing in metazoans. Cell Rep. 42, 112112 (2023).

16. Irimia, M. et al. Evolutionarily conserved A-to-I editing increases protein stability of the alternative splicing factor Nova1. RNA Biol. 9, 12–21 (2012).

17. Zhang, L., Yang, C.-S., Varelas, X. & Monti, S. Altered RNA editing in 3’ UTR perturbs microRNA-mediated regulation of oncogenes and tumor-suppressors. Sci. Rep. 6, 23226 (2016).

18. Park, E., Jiang, Y., Hao, L., Hui, J. & Xing, Y. Genetic variation and microRNA targeting of A-to-I RNA editing fine tune human tissue transcriptomes. Genome Biol. 22, 77 (2021).

19. Salter, J. D., Bennett, R. P. & Smith, H. C. The APOBEC protein family: United by structure, divergent in function. Trends Biochem. Sci. 41, 578–594 (2016).

20. Sharma, S. et al. APOBEC3A cytidine deaminase induces RNA editing in monocytes and macrophages. Nat. Commun. 6, 6881 (2015).

21. Sharma, S. et al. Mitochondrial hypoxic stress induces widespread RNA editing by APOBEC3G in natural killer cells. Genome Biol. 20, 37 (2019).

22. Sharma, S., Patnaik, S. K., Taggart, R. T. & Baysal, B. E. The double-domain cytidine deaminase APOBEC3G is a cellular site-specific RNA editing enzyme. Sci. Rep. 6, 39100 (2016).

23. Bhakta, S. & Tsukahara, T. C-to-U RNA Editing: A Site Directed RNA Editing Tool for Restoration of Genetic Code. Genes 13, (2022).

24. Baysal, B. E. A recurrent stop-codon mutation in succinate dehydrogenase subunit B gene in normal peripheral blood and childhood T-cell acute leukemia. PLoS One 2, e436 (2007).

25. Alqassim, E. Y. et al. RNA editing enzyme APOBEC3A promotes pro-inflammatory M1 macrophage polarization. Commun. Biol. 4, 102 (2021).

26. Sharma, S., Wang, J., Cortes Gomez, E., Taggart, R. T. & Baysal, B. E. Mitochondrial complex II regulates a distinct oxygen sensing mechanism in monocytes. Hum. Mol. Genet. 26, 1328–1339 (2017).

27. Baysal, B. E. et al. Hypoxia-inducible C-to-U coding RNA editing downregulates SDHB in monocytes. PeerJ 1, e152 (2013).

28. Cheng, Q. et al. Sequential conditioning-stimulation reveals distinct gene- and stimulus-specific effects of Type I and II IFN on human macrophage functions. Sci. Rep. 9, 5288 (2019).

29. Navarro, E., et al. G2019S variant is associated with transcriptional changes in Parkinson’s disease human myeloid cells under proinflammatory environment. bioRxiv (2024) doi:10.1101/2024.05.27.594821.

30. Alasoo, K. et al. Transcriptional profiling of macrophages derived from monocytes and iPS cells identifies a conserved response to LPS and novel alternative transcription. Sci. Rep. 5, 12524 (2015).

31. Snijders, G. et al. The human microglia responsome: a resource to better understand microglia states in health and disease. Research Square (2023) doi:10.21203/rs.3.rs-3433713/v1.

32. Picardi, E., D’Erchia, A. M., Lo Giudice, C. & Pesole, G. REDIportal: a comprehensive database of A-to-I RNA editing events in humans. Nucleic Acids Res. 45, D750–D757 (2017).

33. Nalls, M. A. et al. Identification of novel risk loci, causal insights, and heritable risk for Parkinson’s disease: a meta-analysis of genome-wide association studies. Lancet Neurol. 18, 1091–1102 (2019).

34. Saade, M., de Souza, G. A., Scavone, C. & Kinoshita, P. F. The Role of GPNMB in Inflammation. Front. Immunol. 12, (2021).

35. Müller, M., Carter, S., Hofer, M. J. & Campbell, I. L. Review: The chemokine receptor CXCR3 and its ligands CXCL9, CXCL10 and CXCL11 in neuroimmunity – a tale of conflict and conundrum. Neuropathol. Appl. Neurobiol. 36, 368–387 (2010).

36. Marro, B. S. et al. Disrupted CXCR2 Signaling in Oligodendroglia Lineage Cells Enhances Myelin Repair in a Viral Model of Multiple Sclerosis. J. Virol. (2019) doi:10.1128/jvi.00240-19.

37. Trotta, E. On the normalization of the minimum free energy of RNAs by sequence length. PLoS One 9, e113380 (2014).

38. Van Norden, M. et al. The implications of APOBEC3-mediated C-to-U RNA editing for human disease. Commun Biol 7, 529 (2024).

39. Hu, X., Zou, Q., Yao, L. & Yang, X. Survey of the binding preferences of RNA-binding proteins to RNA editing events. Genome Biol. 23, 169 (2022).

40. Chen, L.-L., DeCerbo, J. N. & Carmichael, G. G. Alu element-mediated gene silencing. EMBO J. 27, 1694–1705 (2008).

41. Fu, T. et al. Massively parallel screen uncovers many rare 3′ UTR variants regulating mRNA abundance of cancer driver genes. Nat. Commun. 15, 1–20 (2024).

42. Tan, B. Z., Huang, H., Lam, R. & Soong, T. W. Dynamic regulation of RNA editing of ion channels and receptors in the mammalian nervous system. Mol. Brain 2, 13 (2009).

43. Moore, S. et al. ADAR2 mislocalization and widespread RNA editing aberrations in C9orf72-mediated ALS/FTD. Acta Neuropathol 138, 49–65 (2019).

44. Ma, Y. et al. Atlas of RNA editing events affecting protein expression in aged and Alzheimer’s disease human brain tissue. Nat. Commun. 12, 7035 (2021).

45. Li, W. et al. Transcriptomic analysis reveals associations of blood-based A-to-I editing with Parkinson’s disease. J Neurol 271, 976–985 (2024).

46. Beylina, A., Langston, R. G., Rosen, D., Reed, X. & Cookson, M. R. Generation of fourteen isogenic cell lines for Parkinson’s disease-associated leucine-rich repeat kinase (LRRK2). Stem Cell Res. 53, 102354 (2021).

47. Dobin, A. et al. STAR: ultrafast universal RNA-seq aligner. Bioinformatics 29, 15–21 (2013).

48. Li, B. & Dewey, C. N. RSEM: accurate transcript quantification from RNA-Seq data with or without a reference genome. BMC Bioinformatics 12, 323 (2011).

49. Li, H. et al. The Sequence Alignment/Map format and SAMtools. Bioinformatics 25, 2078–2079 (2009).

50. Piechotta, M., Naarmann-de Vries, I. S., Wang, Q., Altmüller, J. & Dieterich, C. RNA modification mapping with JACUSA2. Genome Biol. 23, 115 (2022).

51. Wang, K., Li, M. & Hakonarson, H. ANNOVAR: functional annotation of genetic variants from high-throughput sequencing data. Nucleic Acids Res. 38, e164 (2010).

52. Hoffman, G. E. & Schadt, E. E. variancePartition: interpreting drivers of variation in complex gene expression studies. BMC Bioinformatics 17, 483 (2016).

53. Ritchie, M. E. et al. limma powers differential expression analyses for RNA-sequencing and microarray studies. Nucleic Acids Res 43, e47 (2015).

54. Robinson, M. D., McCarthy, D. J. & Smyth, G. K. edgeR: a Bioconductor package for differential expression analysis of digital gene expression data. Bioinformatics 26, 139–140 (2010).

55. Law, C. W., Chen, Y., Shi, W. & Smyth, G. K. voom: Precision weights unlock linear model analysis tools for RNA-seq read counts. Genome Biol. 15, R29 (2014).

56. Hofacker, I. L. Vienna RNA secondary structure server. Nucleic Acids Res. 31, (2003).

57. Choudhury, M. et al. Widespread RNA hypoediting in schizophrenia and its relevance to mitochondrial function. Sci Adv 9, eade9997 (2023).

58. Ashuach, T. et al. MPRAnalyze: statistical framework for massively parallel reporter assays. Genome Biol. 20, 183 (2019).

